# Ecology reshapes interactions between antibiotic stressors through metabolic feedbacks

**DOI:** 10.64898/2026.10.21.760830

**Authors:** Xianyi Xiong, Tong Zhou, Ave T. Bisesi, Tsz Fung Wong, Jeremy M. Chacón, Hannah Furseth, Lauren Slavic, Paul C. Blainey, William R. Harcombe

## Abstract

Stressors can combine to inhibit organismal growth or function more (synergy) or less (antagonism) strongly than expected from their individual effects, yet whether these interactions are intrinsic properties of cells or stressors, or emergent consequences of ecological interactions remains unclear. Antibiotic combinations provide a tractable model for studying stressor interactions, but have been characterized largely in bacterial monocultures. Here, we compared antibiotic stressor interactions in *Escherichia coli* growth in monoculture versus when cross-feeding with *Salmonella enterica*. Across 36 antibiotic pairs, cross-feeding frequently shifts interactions toward antagonism and away from synergy. In the sulfamethoxazole-trimethoprim synergy, dual treatment increases carbon excretion by *E. coli,* promoting partner growth and attenuating stressor synergy through reciprocal metabolic feedback. Perturbing partner metabolite usage restores monoculture-like interaction outcomes across multiple drug pairs. Together, our results demonstrate that stressor interactions can emerge from ecological feedbacks mediated by metabolic excretion.

## Introduction

Living organisms, from animals to microbes, are frequently exposed to multiple stressors at the same time^1,2^. Yet responses to combined stressors are difficult to predict from single-stressor effects because stressors can interact to produce outcomes that are stronger or weaker than expected from independence, commonly described as synergy and antagonism, respectively^3–5^. Understanding such interactions is important across biology, from forecasting ecological responses to environmental change to designing combination therapies against disease^1,3,6,7^.

A key challenge is predicting how stressors interact in the ecological contexts in which organisms live. Stressor interactions are often treated as intrinsic properties of the stressors or of the focal organism experiencing them. In natural settings, however, organisms are embedded in ecological networks shaped by competition, facilitation, and mutualism, all of which can alter how stressors propagate through biological systems^8,9^. Work in macroecology has shown that outcomes of stressor interactions can shift across ecological contexts^5,10^, but a mechanistic framework is still missing for predicting when interactions measured in isolation will be preserved, weakened, strengthened or altered in communities. A central unsolved question is therefore whether stressor interactions can emerge from ecological interaction among species, rather than from the stressors or the single species alone.

Synthetic microbial communities offer a tractable setting to answer this question because ecological interactions can be experimentally controlled and linked molecularly to mechanisms. Most current understanding of stressor interactions comes from studying pairs of antibiotic stressors in bacterial monocultures^6,11^. In these studies, drug interactions are typically inferred from changes in growth rate^12^ at sublethal concentrations as an integrative physiological consequence of target inhibition^13^. This framework has yielded important generalities, including the observation that antibiotic stressors targeting related cellular processes often produce similar interaction profiles, whereas ones targeting distinct processes often antagonize one another^13,14^. Genetic studies have further identified intracellular determinants that shape interactions between antibiotic stressors^15,16^, although phylogenetic relatedness cannot predict these interactions^17^. Together, this body of work has provided a powerful monoculture framework for predicting interactions between antibiotic stressors, yet an equivalent framework is missing for predicting how such interactions change when the same species grows in a community.

This missing framework is necessary because interacting species can directly alter each other’s susceptibility and survival especially under antibiotic stressors. Community members may buffer antibiotics, amplify their effects, or indirectly suppress a focal species by disrupting a metabolically interdependent partner^18–21^. Drug stressors can also reshape ecology more subtly by altering the metabolites that microbes release into their environment^22,23^. Because metabolic exchanges determine how species affect one another, antibiotic-induced changes in extracellular metabolism provide a plausible route by which community context could alter antibiotic interactions. Although extracellular metabolic reconfiguration is increasingly recognized as a driver of community responses to single antibiotics^22,24^, whether it can systematically reshape interactions between antibiotic stressors remains unknown.

Addressing this question has been challenging for two reasons. First, ecological effects of single stressors are already difficult to infer from monoculture data^5,23^, making it hard to separate direct drug effects from community-mediated ones. Second, most stressor interaction studies in communities have lacked well-matched monoculture controls^25–27^ or species-resolved measurements^17^. As a result, when interaction outcomes differ in communities, it is often unclear which species experience the shift and what mechanism underlies it^28^. More broadly, this has limited efforts to connect stressor interaction biology with microbial ecology on a mechanistic level.

Here we used a reductionist two-species system to test whether ecology can shape interactions between antibiotic stressors. Cross-feeding is a common ecological interaction in which species of bacteria exchange nutrients to grow in natural environments including soil, freshwater, and even the human gut^9^. We compared antibiotic interactions in *Escherichia coli* grown in supplemented monoculture versus in an obligate cross-feeding mutualism with *Salmonella enterica*^29–30^. Our system allows the physiology of *E. coli* to be closely matched across ecological conditions while selectively adding or removing ecological feedback, and therefore serves as a well-controlled model to answer our question.

Across 36 diverse antibiotic pairs, we found that cross-feeding systematically alters the antibiotic interaction landscape experienced by *E. coli*, with many interactions shifting away from synergy and toward antagonism. Using metabolic modeling, metabolite assays, and genetic perturbations, we showed that these shifts arise because paired antibiotic stress can reconfigure extracellular metabolite excretion, which changes partner growth and feeds back on the focal species. Our results establish a mechanistic framework for studying community-dependent stressor interactions and show that antibiotic interactions can be reshaped by ecological feedbacks mediated through extracellular metabolism.

## Results

### Drug combinations have different effects in monoculture and cross-feeding *E. coli*

To study whether cross-feeding affects interactions between antibiotics, we measured the growth of *E. coli* when cross-feeding with *S. enterica* versus when growing in monoculture. The co-culture is grown in lactose minimal media in which the methionine-auxotrophic *E. coli* excretes carbon (galactose and acetate) that is consumed by *S. enterica*, and in return relies on methionine released by *S. enterica*^29–31^ (Fig. 1A and S1A). *E. coli* is grown under highly similar conditions in monoculture by supplementing the lactose minimal medium with excess methionine (Fig. S1B-E; see Supplementary Text 1 for quantitative support).

**Figure 1.**
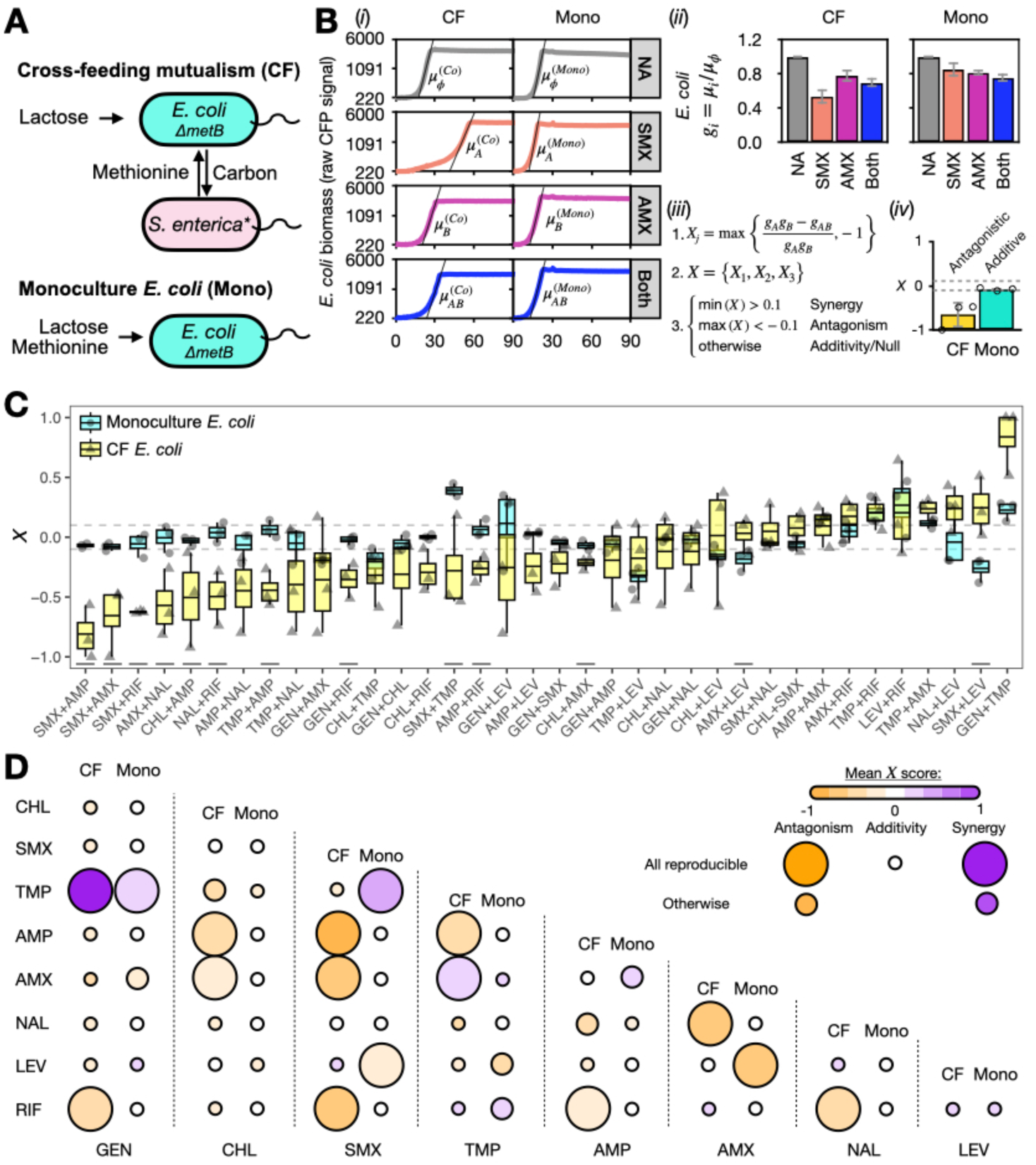
Cross-feeding frequently reduces antibiotic synergy in *E. coli*. **A** Schematic of the cross-feeding mutualism between *ΔmetB E. coli* and methionine-overproducing *S. enterica*, compared with *E. coli* monoculture. Panel adapted from Fig. 1A in Xiong et al.^18^ **B** Measurement of antibiotic interactions. (*i*) Growth rates (*μ*) are determined for *E. coli* grown with no drug, single, sub-MIC drugs, or drug pairs, using log-linear fits (lines) of CFP signals over time. (*ii*) Relative growth rates (*g*) are calculated by normalizing *μ* values to the no-drug control (*μ_φ_*). Means ± SD from three independent experiments are shown. (*iii-iv*) Antibiotic interaction indices (*X*) are computed per experiment *j* and averaged. Interaction type (synergistic, antagonistic, or additive/null) is defined based on thresholds. **C** Antibiotic interaction indices (*X*) across three biological replicates for monoculture (Mono) and cross-feeding (CF) *E. coli*. Grey dashed lines denote *X =* 0.1 and -0.1. Horizontal lines denote drug pairs with different antibiotic interaction classifications as well as reproducibly higher *X* scores in either monoculture or cross-feeding *E. coli*. Antibiotic abbreviations are listed as in Fig. S2A. **D** Predicted interaction types for monoculture (Mono) and cross-feeding (CF) *E. coli*. “All reproducible” refers to classified synergistic or antagonistic interactions with reproducibly higher or lower *X* scores in the classified ecological condition than the opposite one.

We used a growth rate-based screen at standardized sub-MIC concentrations of diverse antibiotics as an established high-throughput method^13^ to ask whether ecological context alters drug interactions at matched exposures. We first chose nine antibacterial drugs from five representative antibiotic classes^32,33^ and measured their MICs in monocultures of *E. coli* in lactose and methionine, and monocultures of *S. enterica* in galactose^31^ (Fig. S1A and S2A-B). We then determined each drug’s working concentration as one sub-MIC dosage that allows growth in all conditions but slows the *E. coli* growth rates to between 60% and 95% of the no-drug control (Fig. S1A and S2C-F). Because cultures were inoculated during exponential growth at low starting densities, the growth curves often show an apparent lag before signal becomes detectable; however, based on the no-drug growth rates, much of this delay is expected from the time required for exponential growth to rise above the instrument detection limit rather than from a true biological lag (Supplementary Text 1). We therefore focused our analysis on growth rates rather than apparent lag times.

The interaction between two antibiotics is determined from the inhibition of *E. coli* growth rates under both drugs compared to each antibiotic’s effect alone (Fig. 1B and S2C). In an example antibiotic pair between sulfamethoxazole (SMX; Drug A) and amoxicillin (AMX; Drug B), we fit a log-linear function to the *E. coli* growth curve in monocultures and co-cultures to obtain average growth rates in each species without antibiotics (*μ_φ_*), with SMX alone (*μ_A_*), with AMX alone (*μ_B_*), and with both SMX and AMX (*μ_AB_*) (Fig. 1B and S2C-F).

To assign antibiotic pairs to interaction types, we followed the Bliss independence model^13,34^. The relative growth rate of each individual drug (*g_i_* = *μ_i_*/*μ_φ_*) was used to calculate a predicted growth rate of the antibiotic combination if the pair acts additively (*g_A_g_B_*) without apparent interactions. We subtracted our observed relative growth rate (*g_AB_* = *μ_AB_*/*μ_φ_*) with both drugs from the predicted relative growth rate to get an antibiotic interaction index (*X*). Conservatively, values of *X* from three independent experiments (different colonies, weeks, and antibiotic stocks) were used to determine the pairwise antibiotic interaction classifications, and then, following previous work^13^, a threshold approach was used for classification. Drug interactions were classified as synergistic if all *X* were greater than 0.1 [min(*X*) > 0.1], antagonistic if all *X* were less than -0.1 [max(*X*) < -0.1], or otherwise additive (aka null interaction) in monocultures and co-cultures, respectively (Fig. 1B; Materials & Methods).

We found that cross-feeding had a substantial impact on drug interactions (Fig. 1C and S3A-B). In 39% (13) of the 36 antibiotic pairs, the antibiotic interaction has different classifications for monoculture versus cross-feeding *E. coli* (Fig. 1D). We also compared the values of *X* across drug types and found that for 26 (72%) of the 36 combinations tested, average *X* scores were more negative for cross-feeding *E. coli* (*P =* 0.015, Binomial test). The observation that more drug pairs shift toward antagonism than toward synergy is also robust across antibiotic interaction classification methods and cutoff values of *X* (Fig. S3C). We further verified that the frequent shifts toward antagonism are not associated with evolved resistance in *E. coli* (Fig. S4).

### The “weakest-link” hypothesis explains a subset of the observed differences in antibiotic interaction

Why do many pairwise antibiotic interactions shift toward antagonism in co-culture? In obligate cross-feeding mutualisms, community growth may be limited by the more susceptible species, or the “weakest link”^21^. Under this framework, if *S. enterica* responds to a drug combination differently from *E. coli* and is the more susceptible partner, then the interaction observed for *E. coli* in co-culture may align with the drug interaction profile of *S. enterica*^35^.

This framework can explain the shift in *E. coli* response to AMX (amoxicillin) and nalidixic acid (NAL). In monoculture, *S. enterica* is fourfold more susceptible to AMX and fortyfold more susceptible to NAL than *E. coli* (Fig. S2A-B), and in co-culture it also grows more slowly than *E. coli* in the presence of each single drug (Fig. S2E). Consistent with the weakest-link expectation, the antagonistic interaction observed for cross-feeding *E. coli* is matched by the antagonistic interaction seen in monoculture *S. enterica* (Fig. S5A-B).

However, the weakest-link hypothesis does not explain most cases in which antibiotic interactions shift toward antagonism in co-culture, nor are these cases explained by the interaction type observed in either monoculture alone (Fig. S5C). Many drug pairs shift in co-culture even when *E. coli* and *S. enterica* have similar susceptibilities, share the same interaction type in monoculture, or when *E. coli* remains the slower-growing species under drug treatment (Figs. S2D-F). Although co-culturing *E. coli* induces smaller relative single-drug growth rates (*g_i_*) than monoculture, this single-drug growth rate difference between mono- and co-culture does not strongly correlate with the mean difference in *X* scores (Fig. S5D). Thus, shifts toward antagonistic interactions between antibiotic stressors in co-culture cannot be understood solely from the most susceptible partner or changes in growth rate alone, suggesting additional ecological mechanisms that shape the co-culture response.

### Models predict that changes in stressor interaction are mainly driven by metabolism

We implemented genome-scale metabolic models to test whether paired stressors also interact differently for cross-feeding *E. coli*. Genome-scale metabolic models can predict growth and metabolic fluxes from cellular reaction networks and have been used to model antibiotic perturbations by reducing the efficiency of drug-targeted reactions^36^. In this work, we extended this framework to a multi-species context using COMETS^30,37^, which simulates growth and metabolic exchange in microbial communities (Fig. 2A and S6A; Materials & Methods).

**Figure 2.**
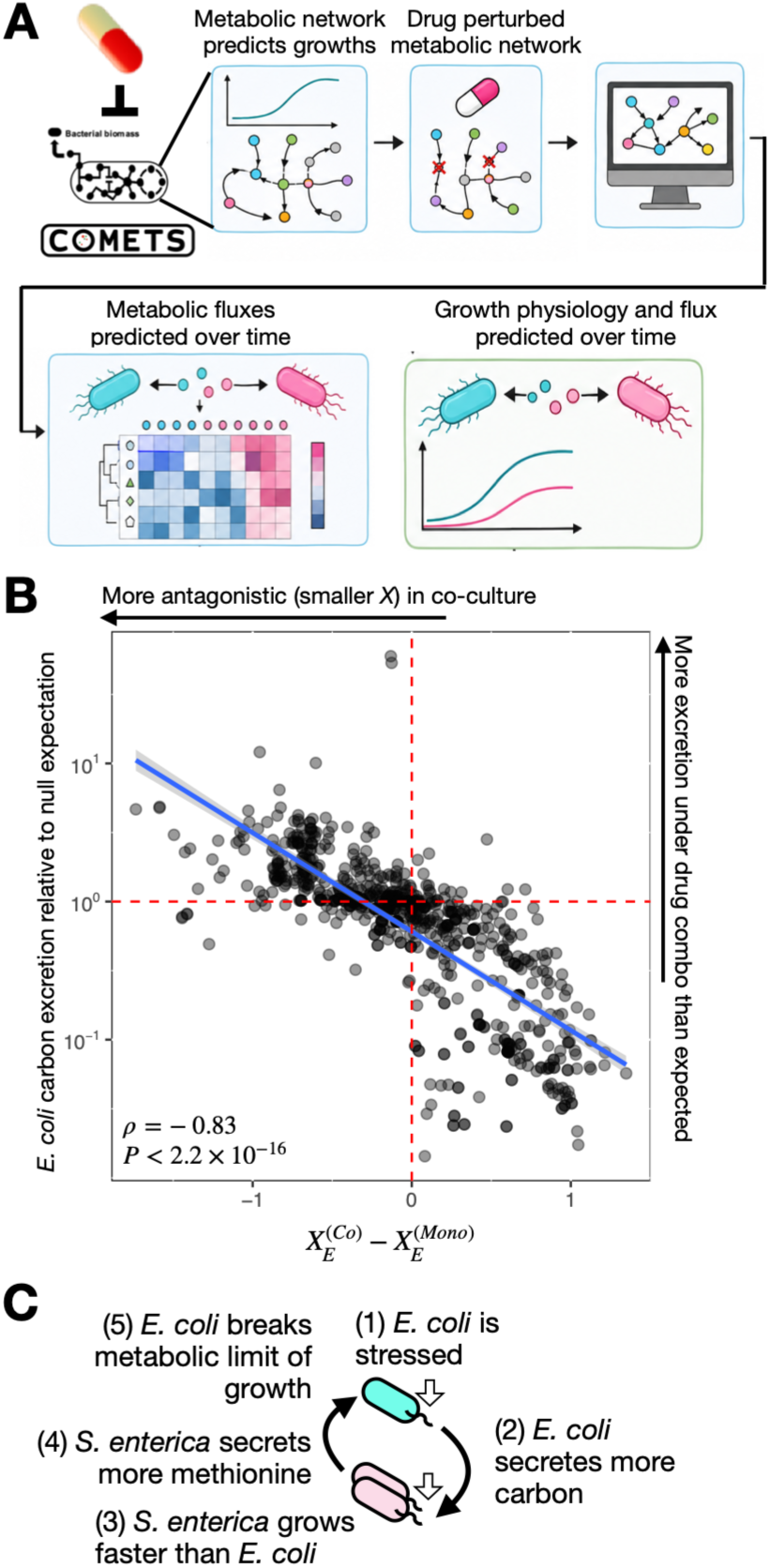
Metabolic models link metabolite excretion to shifts toward antagonistic stressor interaction for cross-feeding *E. coli*. **A** Schematic for the metabolic models on pairwise gene perturbation for monoculture and cross-feeding *E. coli*. **B** Differences in the stressor interaction indices (*X*) between cross-feeding (Co) and monoculture (Mono) *E. coli* (*E*) are negatively correlated with the total carbon excretion (total carbon from galactose and acetate excretions) flux from *E. coli* under dual perturbation relative to the null expectation from each single perturbation. Vertical dashed line indicates no change in stressor interaction. Horizontal dashed line indicates carbon excretion in dual perturbation matches null expectation. **C** Schematic of the proposed model explaining weakened synergy and increased antagonism in cross-feeding *E. coli*.

Critically, we were not attempting to model the mechanisms of action for the antibiotics in our experiment, but rather we aimed to use the models to investigate the ecological effects of combined perturbations generally. Altering the efficiency of metabolic reactions has previously been shown to recapitulate observed ranges of drug interactions from antagonism to synergy^36^. Thus, this approach allows us to explore a diverse set of perturbations that many alternative modeling approaches cannot easily simulate.

We broadly perturbed 40 gene targets carrying out different metabolic reactions and simulated the effects of inhibiting these genes individually and in all 780 pairs for monoculture and cross-feeding *E. coli* (Fig. S6B-E). These simulations align with the antibiotic screen in that the interaction types of perturbing many gene pairs also differ with and without cross-feeding (Fig. S6C-D and Supplementary Text 2). Our models also suggest that changes in growth rate alone may not be the sole reason for the observed shifts in stressor interaction (Fig. S7A and Supplementary Text 2).

Instead, the models frequently reveal non-additive effects of pairwise stressors on extracellular metabolic fluxes, particularly carbon excretion by *E. coli* in cross-feeding co-culture (Fig. S7B-C). Bigger shifts toward stressor antagonism in co-culture are associated with more *E. coli* carbon excretion under stressor combinations than expected from the additive predictions under single stressors (Fig. 2B). Consistent with this result, we also observed a modest experimental trend for antibiotic stressor combinations that shift toward antagonism in co-culture to produce higher final *S. enterica* frequencies (Fig. S7D).

Based on these results, we hypothesize that extracellular metabolic reconfiguration is an ecological mechanism by which cross-feeding shifts antibiotic interactions toward antagonism and away from synergy (Fig. 2C). Increased antagonism between drugs may be observed because the drugs drive increased carbon excretion in *E. coli* (Fig. S7B-C). This increased carbon excretion by *E. coli* will increase the resources available for *S. enterica* and drive an increase in *S. enterica* growth (Fig. S7D). This increased *S. enterica* growth then increases available methionine^38^, generating feedback that reduces the metabolic constraints on *E. coli* in co-culture.

### Carbon excretion breaks SMX-TMP synergy for cross-feeding *E. coli*

The sulfamethoxazole-trimethoprim (SMX-TMP) synergy in monoculture *E. coli* arises from the two drugs targeting sequential reactions in folate biosynthesis controlled by *folP* and *folA*, respectively^33,39^, so this drug pair is also convenient for metabolic simulations. Weakened SMX-TMP synergy in cross-feeding *E. coli* compared with monoculture was consistently observed across checkerboard assays and additional growth curve experiments (Fig. S8A-D). In monocultures, *S. enterica* and *E. coli* are equally susceptible to SMX and TMP, respectively (Fig. S2B). In co-cultures, *S. enterica* never grows faster (*μ_i_*) than *E. coli* without drugs or with either SMX or TMP, regardless of whether *S. enterica* grows by obligately cross-feeding with *E. coli* or through unidirectional facilitation by carbon-providing *E. coli* (Fig. S2E and S8E). In contrast, when both SMX and TMP are present in either cross-feeding or facilitative co-cultures, *S. enterica* grows significantly faster than *E. coli* and reaches higher final frequencies (Fig. 3A-B and S8F-I). These patterns are consistent with the SMX-TMP combination driving a substantial change in *E. coli* excretion.

**Figure 3.**
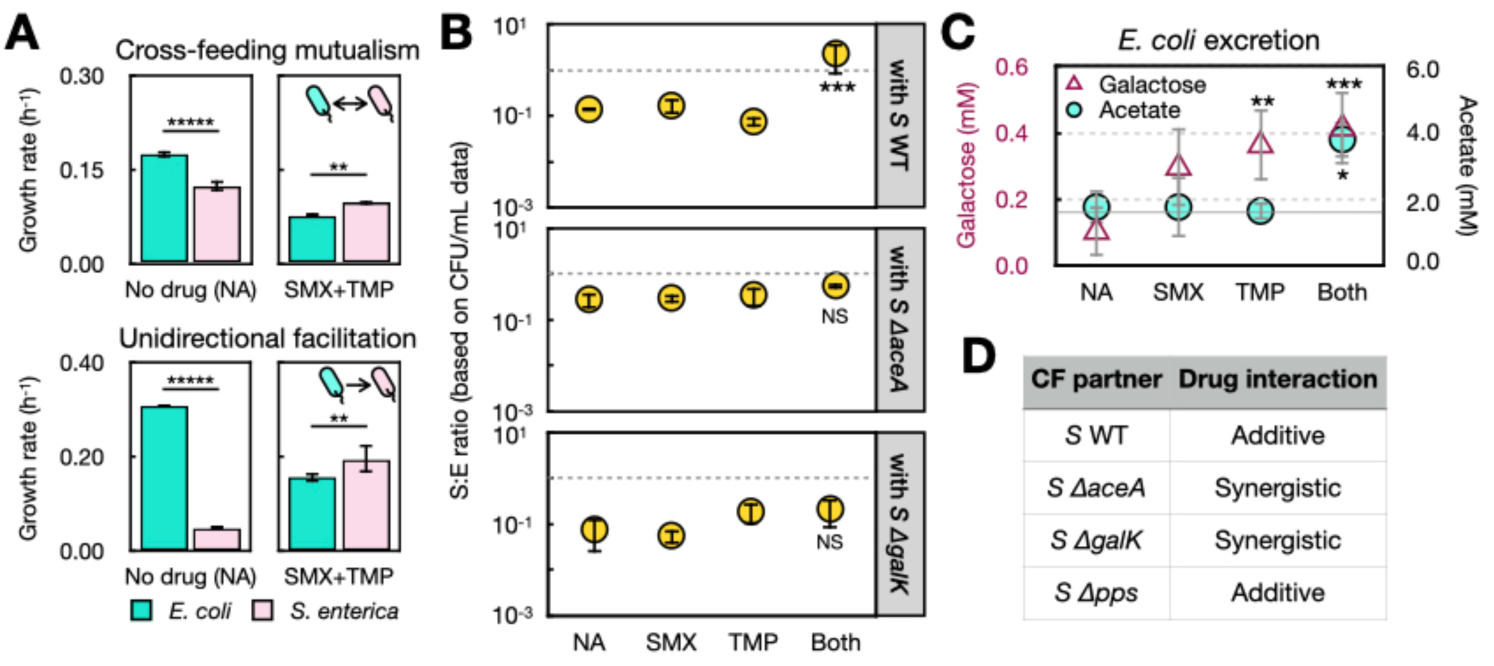
Extracellular metabolic reconfiguration weakens SMX-TMP synergy in cross-feeding *E. coli*. **A** Raw growth rates (*μ_i_*) of both species with and without both drugs in the cross-feeding vs the unidirectional facilitative co-culture. In the facilitative co-culture, *S. enterica* is metabolically dependent on *E. coli* but not vice versa. **B** Ratios between S:E CFU/mL counts from selective plating of co-culturing *E. coli* with wild-type (WT), *ΔaceA*, or *ΔgalK S. enterica* (*S*) in the cross-feeding mutualism. Marks of statistical comparisons are from comparing each antibiotic condition against the NA counterpart. **C** Acetate (circle) and galactose (triangle) concentrations measured from log-phase (OD_600_=0.21) monoculture *E. coli* during growth under different antibiotic conditions. Dashed line is the expected acetate production under SMX+TMP (both) based on the null/additive expectation from SMX and TMP alone. Marks of statistical comparisons are from comparing each antibiotic condition against the NA counterpart. **D** Antibiotic interaction for *E. coli* when cross-feeding with various *S. enterica* strains. Statistical tests are done with one-way ANOVA tests, and Tukey’s HSD tests are run for all panels of figures with more than two groups of data. NS: *P* ≥ 0.1; *: 0.01 ≤ *P* < 0.05; **: 0.001 ≤ *P* < 0.01. ***: 0.0001 ≤ *P* < 0.001. ****: 0.00001 ≤ *P* < 0.0001. These conventions are used throughout the manuscript.

We next found evidence of changes in excretion and the effect on weakening SMX-TMP synergy (Fig. 3C and S9). Both metabolic models and experiments agree that SMX and TMP together increase *E. coli* excretion of key metabolites, acetate and galactose, on which *S. enterica* relies for growth. Under both drugs, acetate is non-additively higher while galactose is simply highest among all conditions (Fig. 3C). Consistent with this result, the co-culture is no longer dominated by *S. enterica* when the *S. enterica* strains cannot metabolize galactose (*ΔgalK*) or acetate (*ΔaceA*), respectively (Fig. 3B and S10A). Cross-feeding with either *ΔgalK* or *ΔaceA S. enterica* is sufficient to drive a synergistic SMX-TMP interaction in *E. coli*—just like its monoculture counterpart (Fig. 3D and S10B). The knockouts do not reduce growth of the co-culture in the absence of antibiotics (Fig. S10C), nor do they alter the resistance of *S. enterica* to either drug (Fig. S10D).

We did not find evidence of other organic carbon metabolites being over-excreted by *E. coli* in the presence of SMX and TMP (Fig. S10E-I). For example, *Δpps S. enterica,* which is unable to metabolize lactate^40^, results in an additive SMX-TMP interaction in cross-feeding *E. coli* just like the wild type (Fig. 3D and S10B). The frequency of the *S. enterica Δpps* mutant also increases when it cross-feeds with *E. coli* under both SMX and TMP (Fig. S10E). Together, our results here support extracellular metabolic reconfiguration as an ecological mechanism that weakens the SMX-TMP synergy (Fig. 2C).

### Metabolic excretion frequently shifts antibiotic interaction toward antagonism in co-culture

We compared fold changes in the antibiotic-induced S:E ratios in cross-feeding co-cultures harboring different genetic backgrounds of *S. enterica* (Fig. 4A). When *E. coli* cross-feeds with wild-type (WT) *S. enterica*, the *S. enterica* frequency also increases in 5 out of 10 drug pairs besides SMX+TMP that also shift toward antibiotic antagonism in co-culture. For each of these 5 pairs, the increase in *S. enterica* frequency could be attributed to at least one antibiotic in the pair, whose single-drug effect increases S:E ratios compared to the no-drug control in the WT background (red stars). This pattern is consistent with *S. enterica* receiving growth benefits from antibiotic-induced *E. coli* excretion.

**Figure 4.**
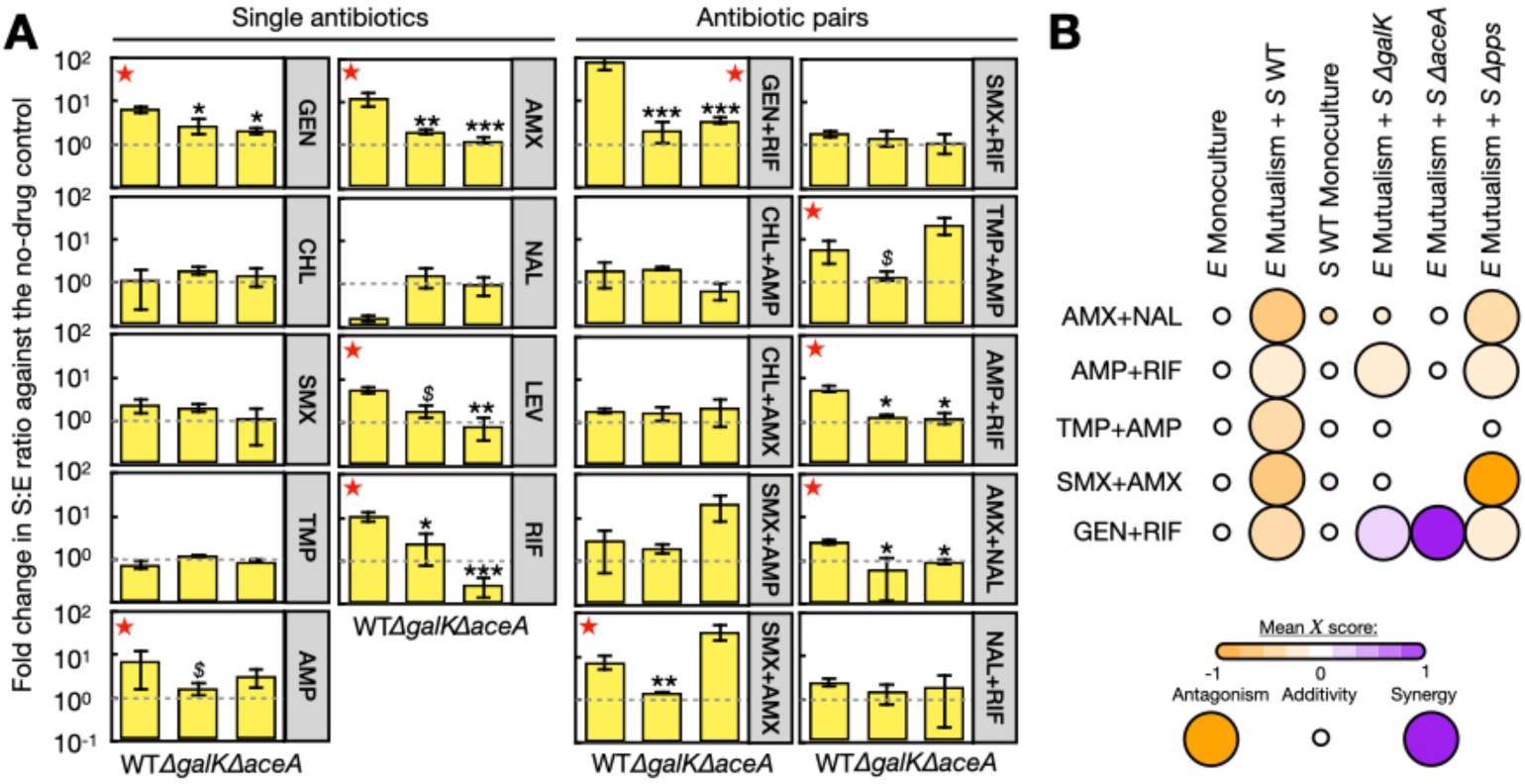
Extracellular metabolic reconfiguration may be a general mechanism driving shifts toward antibiotic antagonism in cross-feeding *E. coli*. **A** Fold changes in S:E ratios based on CFU counts across antibiotic treatments relative to the no-drug control in cross-feeding communities with *S. enterica* WT, *ΔgalK*, and *ΔaceA* strains, respectively. Red stars: antibiotic treatments where disrupting galactose or acetate metabolism in *S. enterica* prevents the antibiotic-induced increase in S:E ratios. *$*: 0.05 ≤ *P* < 0.1 with Tukey’ HSD. **B** Antibiotic interaction types in *E. coli* (*E*) for the five drug pairs marked by red stars in **A**. Graphic demonstration of antibiotic interaction follows Fig. 1D.

For all five pairs and for each implicated single antibiotic (red stars), replacing WT *S. enterica* with either the *ΔgalK* or *ΔaceA* mutant is sufficient to abolish the increase in *S. enterica* frequencies (Fig. 4A). Cross-feeding with either *S. enterica ΔgalK* or *ΔaceA* also usually breaks the antibiotic antagonism experienced by *E. coli* (Fig. 4B and S10J). Together, these results suggest that extracellular metabolic reconfiguration is a common mechanism by which cross-feeding shifts antibiotic interactions towards antagonism in *E. coli* (Fig. 1D and 2C).

## Discussion

In this study, we showed that antibiotic interactions are not fixed properties of drugs acting on isolated cells but instead can vary with ecological context (Fig. 1C-D). By comparing *E. coli* in supplemented monoculture versus obligate cross-feeding co-culture, we found that many antibiotic pairs shift toward lower antibiotic interaction indices (*X*) in co-culture, weakening synergy and strengthening antagonism (Fig. 1C-D and S3B). This result suggests that drug interactions measured in monoculture do not always predict how the same bacteria will respond in a community, because community contexts within which stressors act also have an important, emergent role on the effect of combined stressors.

A central finding of this study is that stressor-induced metabolic excretion can reshape interactions between antibiotic stressors (Fig. 2-4). For example, in the SMX-TMP antibiotic pair, dual drug treatment causes *E. coli* to excrete more acetate and galactose, thereby changing the resources it provides to its partner and promoting *S. enterica* growth^36^, which in turn weakens the expected synergy (Fig. 3). This effect cannot be explained by drug resistance or by growth-rate differences when cross-feeding with different *S. enterica* mutants (Fig. S8 and S10C-D), and it was consistent with the model prediction that paired perturbations can alter metabolite exchange non-additively (Fig. 2B). More broadly, disrupting galactose or acetate utilization in *S. enterica* weakens antibiotic antagonism in *E. coli* across multiple drug pairs beyond SMX and TMP (Fig. 4), indicating that stress-induced changes in an environmentally expressed metabolic trait—rather than co-culture alone—underlies much of the effect.

Our findings also refine the scope of the weakest-link hypothesis. The weakest-link framework provides a useful null expectation when antibiotics alter growth inhibition without substantially changing metabolic exchange, in which case co-culture outcomes may be determined by the more susceptible species. We observed examples that are consistent with this expectation, but weakest-link logic alone cannot explain most drug pairs that shift toward antagonism in co-culture (Fig. S5C). Instead, our data support a broader model in which antibiotics change the ecological function of the community, thereby altering partner benefits and reshaping which species is limiting community growth (Figs. 2-4). Even the AMX-NAL pair, which appears initially consistent with weakest-link expectations, shows an increase in the *S. enterica* frequency under AMX-containing conditions that is more consistent with our metabolic feedback model (Fig. 4 and S5A). Moreover, perturbing partner metabolite usage is sufficient to reduce *S. enterica* dominance and restore a more monoculture-like interaction outcome (Fig. 4). Thus, while the weakest-link hypothesis captures cases in which ecology passively transmits stress, our results show *how* stressor combinations can also actively reshape ecology to affect stressor responses.

Although our reductionist approach makes it possible to isolate the ecological mechanism, a few caveats remain. Our study relies on a simplified two-species system and a screening design based on one sub-MIC concentration for each antibiotic. The alteration of metabolite exchange is likely not the only mechanism that contributes to antibiotic interactions being different in monoculture and co-culture. Our models that simulate genetic perturbations at best mimic the effects of metabolism-targeting antibiotics using only one mode of growth constraints in the flux balance analysis that the COMETS platform relies on. Our models also do not capture the diverse modes of action of the other antibiotics that do not work by targeting bacterial metabolism. Future work should identify the molecular mechanisms by which antibiotics alter extracellular metabolite excretion and explain why some remaining drug pairs switch interaction classifications without changing their species ratios.

In summary, we showed that extracellular metabolic reconfiguration could act as an ecological driver of altered antibiotic interactions in microbial communities. These findings challenge the monoculture-centric view of antibiotic stressor interaction studies and suggest that responses to antibiotic combinations may depend in part on surrounding microbial partners, because antibiotics can change not only bacterial growth, but also the functional traits through which bacteria affect one another.

## Materials and Methods

### Bacterial strains and study system

*Escherichia coli* used is a Keio line (JW3910) with a *metB* deletion^41^, a *lac* operon reinserted into its genome^29^, and a genome-encoded CFP gene^30^. *S. enterica* (referred to as “*S. enterica* WT”) is a methionine-overproducer strain evolved from *S. enterica* serovar Typhimurium LT2^29^. It encodes a base-pair change in *metA*^38^, an insertion sequence element before *metJ*^42^, and a YFP gene that is later inserted into its genome^30^. *S. enterica ΔaceA* and *ΔilvA* mutants were acquired from previous work^31^; they were constructed by inserting a kanamycin resistant (KanR) cassette in the gene of interest via P22 transduction, and were demonstrated to have expected loss-of-function phenotypes. The *ΔgalK* and *Δpps* mutants were made in this study following the same protocol as in Martinson et al.^31^ All mutants were created in the genetic background without the YFP gene, and the mutants barely affect the co-culture growth physiology without drugs (Fig. S10C). *S. enterica* 14028S WT, and its *ΔydhC* and *Δppc* mutants (Biodefense and Emerging Infections Repository) were also used in this study.

The two-species system composed of *E. coli* and the *S. enterica* methionine overproducer can toggle between cross-feeding mutualism and independent growth. In lactose minimal medium, *E. coli* metabolizes lactose to feed the methionine-overproducing *S. enterica* carbon (galactose and acetate) and obtains methionine from the latter. *E. coli* (*S. enterica*) can be grown independently by supplementing lactose and methionine (galactose) in monocultures. A competitive co-culture between *E. coli* and *S. enterica* can be studied in Hypho media supplemented with glucose and methionine. A unidirectional facilitative co-culture between the two species is studied in the Hypho media with lactose and methionine, where *S. enterica* relies on *E. coli* for carbon but *E. coli* can independently obtain nutrient from the media.

### Media conditions and growth experiments

Growth experiments were performed in Hypho minimal medium or on minimal agar. All agar plates contain 1% (m/v) agar. Hypho medium^43^ is a buffered medium that contains 14.52 mM K_2_HPO_4_, 18.76 mM NaH_2_PO_4_, 3.78 mM (NH_4_)_2_SO_4_, 0.82 mM MgSO_4_, 0.6 μM ZnSO_4_, 9.98 μM CaCl_2_, 0.5 μM MnCl_2_, 1 μM (NH_4_)_6_Mo_7_, 0.5 μM CuSO_4_, 1 μM CoCl_2_, 0.169 μM NaWO_4_, and 8.88 μM FeSO_2_. The cross-feeding media is supplemented with 2.82 mM lactose, whereas the *E. coli* monoculture media with 2.82 mM lactose and 80 μM methionine. The *S. enterica* monoculture media is supplemented with 5.56 mM galactose or glucose, or 8.46 mM succinate, whenever noted.

A colony of every strain was grown in LB media till early log phase (OD_600_ = 0.2∼0.4) in 5 mL cultures with shaking at 200 rpm at 37°C. Cells were then washed three times in and diluted into 200 μL fresh Hypho media to start growth experiments at OD_600_ = 0.001 in 96-well plates, which were incubated with shaking at 485 rpm at 37°C for up to one week till cultures reach the stationary phase, unless otherwise noted. Parafilm was taped around the edge of each 96-well plate to reduce water loss without depleting oxygen. The low starting OD_600_ was deployed in the growth experiment to minimize spontaneous mutations following convention.

Growth dynamics was measured on a Tecan InfinitePro 200 plate reader (Tecan US, Inc., NC). At every 20 min for up to six days, OD_600_, CFP (Ex: 430 nm, Em: 480 nm, for *E. coli*) and YFP (Ex: 500 nm; Em: 530 nm, for *S. enterica*) were recorded for every well in the 96-well plate. Wells on the outer rim were not used due to edge effects and instead contain 200 μL autoclaved water to reduce water loss in experimental wells. Every experiment was repeated in at least three biological replicates. Final densities of each species were assayed as the end-of-experiment fluorescent protein signals or colony forming units (CFUs) on selective agar. Hypho minimal agar with lactose + methionine or with galactose was used to selectively grow *E. coli* and *S. enterica*, respectively, for 48 hours. Every spot of 5 μL from 10^-1^∼10^-8^ dilutions was plated on agar, and spots both at the fewest dilution and are with 10∼50 colonies were counted to calculate species-specific population sizes. For *S. enterica ΔgalK* and 14028S *Δppc*, Hypho agar with glucose or succinate were used for plating, respectively.

We fit a custom-built log-linear curve from our previous work^18^ to growth curves of fluorescence data. For *S. enterica* mutants that were constructed in a background without YFP, growth curves are made from OD_600_ data in monoculture. Growth signals can drop towards the end of the experiment as water loss eventually happens, which prevents reliable curve fit. Thus, because we only care about growth rates, the highest growth signal in the growth curve was manually used to override all signals following its time point. The log-linear equation is:

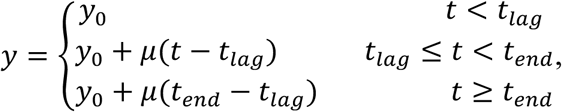

where *y* and *y*_0_ are the natural logarithms of the species abundances at *t* and *t*_0_, respectively. Lag time and the time of end of growth are denoted by *t*_lag_ and *t*_end_, respectively. Growth rate (*μ*) is the slope of the exponential phase of the growth curve. Because log-phase cultures were inoculated below the detection limit of the plate reader, early apparent lag largely reflects the time required for exponential growth to reach detectable OD_600_. Under antibiotic treatment, additional apparent lag may arise from physiological adaptation and reduced net growth rate. Therefore, growth rates were estimated from the log-linear region after cultures exceeded the detection threshold. Bacteria have doubled for ∼6.6 times before the cell density reaches the lower bound of the detection limit and the plate reader begins to measure reliable growth yields over time (Supplementary Text 1).

### Estimation of antibiotic resistance

Nine antibiotics were used in this study: gentamicin (GEN), chloramphenicol (CHL), sulfamethoxazole (SMX), trimethoprim (TMP), ampicillin (AMP), amoxicillin (AMX), nalidixic acid (NAL), levofloxacin (LEV), and rifampicin (RIF) (Fig. S2). Antibiotic resistance in *E. coli* and *S. enterica* strain stocks were measured as the minimum inhibitory concentration (MIC) of each antibiotic that can inhibit full growth in 48 hours in monoculture Hypho media. Antibiotics were diluted in Hypho media to 100 μg/mL. A concentration gradient was created by 2-fold serial dilution in 96-well plates. The lowest concentration that results in no visible turbidity to the naked eye was considered the MIC, which is equivalent to resulting an OD_600_ of <10% of the no-drug counterpart (i.e. MIC_90_). When species-specific MICs are being measured in growing co-cultures, growth is defined as present if the FLP signal or the natural logarithm of CFU counts reaches 10% of the no-drug counterpart.

Our previously established protocol^18^ was used to determine whether resistant mutants appear in co-culture experiments. To estimate *E. coli* (*S. enterica*) resistance, cells from the end of growth are diluted to OD_600_ = 0.001 for monocultures, and OD_600_ = 0.002 for co-cultures, respectively. Cells are resuspended in Hypho media supplemented with lactose and methionine or with galactose before going through the same MIC measurement protocol above. For co-culture cells in these media conditions, the focal species will be able to grow independently, but the partner species will still depend on the focal one. So even though the partner species has a higher monoculture MIC, no growth will occur due to the partner species’ metabolic dependency^18,21^.

### Determination of antibiotic interactions

We assessed the bacterial growth rates during log phase after seven doubles in drug-free media, and in media with single or pairs of drugs at matched sublethal concentrations (Methods & Supplementary Text 1). Interactions between antibiotics are measured using growth rate-based dose-response curves^13^ and yield-based checkerboard assays^44^. For the growth rate method, the antibiotic interaction index (*X*) is calculated in the following equation:

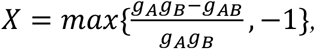

where *g_A_*, *g_B_*, and *g_AB_* are growth rates under drug *A*, drug *B*, and both drugs *A* and *B*, that are normalized to the growth rate in the no-drug condition. Note that the equation limits *X* to be between -1 and 1. Synergistic [min(*X*) > 0.1] and antagonistic [max(*X*) < -0.1] interaction types are classified based on extreme values of *X.* We classified an antibiotic stressor pair to have no interaction (none) when neither of the two criteria are met.

The growth yield-based method was used to confirm interactions between SMX and TMP following an established protocol^33,44^. Briefly, gradients of both SMX and TMP were created in a total of 36 or 64 wells in a 96-well plate, with the highest well matching each drug’s expected MIC. Final growth was read once the no-drug control has entered the stationary phase (2 days for monoculture, 4 days for cross-feeding co-culture). For each edge well without growth, the fractional inhibitory concentration index (FICI) was calculated as follows:

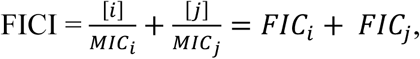

where *FIC_i_* and *FIC_j_* are fractional inhibitory concentrations for SMX and TMP, respectively. A small minimal FICI value (FICI_m_) from all FICI values (FICI_m_ ≤ 0.5) suggests a synergistic interaction, and 0.5<FICI_m_≤4 suggests no explicit interaction is observed^43^.

### Measurement of *E. coli* galactose and acetate excretions

Metabolic excretion was measured in the spent media of monoculture *E. coli*. *E. coli* is grown in 5 mL Hypho lactose and methionine media with no drugs, SMX, TMP, and both SMX and TMP, respectively, till OD_600_ = 0.21. All spent media samples are stored at 4°C in light-protective boxes when not in use. Samples are analyzed within one week of collection.

Acetate excretion was measured by high-performance chromatography (HPLC) assays in 100 μL volumes. The spent media were run on an Agilent 1100 HPLC system with a ThermoFisher Acclaim Mixed-Mode WAX-1 column (4.6 × 150 mm, 5 µm particle size) at 30 °C, flow 0.8 mL/min, mobile phase 25 mmol phosphate buffer (pH 6.0), and monitored with a diode array detector at 210 nm. Chemical signals (mAU) were used to calculate the area under the curve (AUC) for the acetate peak during 7∼9 min retention times. A standard curve was made by adding acetate up to 125 mM in Hypho lactose and methionine media (Fig. S9B).

Galactose excretion was not measured with HPLC because the metal peak in the Hypho media masks the galactose peak. Instead it was measured by comparing differential growth in wild-type and *ΔgalK S. enterica*. These two *S. enterica* strains were grown in spent media separately for 7 days to calculate the difference in final growth yields (ΔOD_600_), which positively correlates with the galactose concentration in the media (Fig. S9C). A standard curve was made by repeating the same growth comparison experiment in Hypho lactose and methionine media with galactose added up to 1.8 mM (Fig. S9C).

### Computational simulation

Simulations were run on the genome-scale metabolic model platform, <u>C</u>omputation of <u>M</u>icrobial <u>E</u>cosystems in <u>T</u>ime and <u>S</u>pace^37^ (COMETS v2.0). In our model, we used flux balance analysis (FBA) to approximate the metabolic impact of antibiotics using the established flux diversion method^36^, which reduces the activity of a single gene, representing inhibition by a single drug. We chose a total of 40 genes (with 780 resultant gene pairs) to perturb. Genes were chosen at random to represent a variety of potential effects, with some genes essential in both *E. coli* and *S. enterica* models, some genes essential in neither, and some genes essential in only one species. A drug was simulated by constraining the stoichiometric coefficients of all metabolic products for each reaction associated with a single gene. Coefficients were constrained to a fraction (1/*α*), such that an irreversible reaction *A* → *B* became 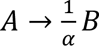 and a reversible reaction *A* → *B* 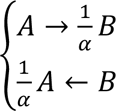. Reactions were constrained even if they could be controlled by another, non-perturbed gene. When evaluating gene pairs, if a reaction is shared between each gene in the antibiotic pair, the corresponding reactants are inhibited according to whichever individual gene imposes the greatest constraint. Shared reactions only exist in six pairs out of the 780 tested. Strain-specific *α* values for each individual tested gene were generated using FBA in CobraPy^44^ v. 0.29.0 and a binary search algorithm to identify a constraint that reduces monoculture growth to 50% to a minimum tolerance of 1E-3 in fewer than 50 search cycles. If constraining a non-essential gene has no impact on growth, the *α* value was set to 100,000. The *α* values vary depending on nutrient conditions and were calculated for *S. enterica* using a minimal media with galactose, while *α* values for *E. coli* were calculated using a methionine-limited minimal media with lactose as the carbon source.

After individual *α* values were calculated using FBA, single or double-drug constraints were then applied to models prior to simulations using dFBA in COMETS v. 0.5.2. The existing *E. coli* model iML1515 with a knockout of the *ΔmetB* gene and a methionine-secreting version of the *S. enterica* model STM_v1_0 were used for all simulations, including for *α* generation. All monoculture and coculture simulations in a liquid batch culture environment were performed until strains reach stationary phase with a timestep of 1 hour. Initial populations sizes were set to 1E-8 grams of dry weight (gDW) for each strain for all simulations and the objective function of each species was set to its core biomass reaction. Lactose serves as the sole carbon source in *E. coli* monoculture and cross-feeding simulations, while galactose was used as the sole carbon source in *S. enterica* monoculture simulations. Methionine is supplemented into the *E. coli* monoculture simulations to enable growth. All non-carbon nutrients not provided during cross-feeding are expected to be present in excess and are supplemented with 1000 mmol per nutrient. Species biomasses and exchange fluxes were recorded and evaluated across the growth period. Log-linear growth rates were estimated using biomass curves as described above and drug interactions were evaluated as before based on those estimates. Optimization was completed with the Gurobi optimizer^45^.

### Statistics

We ran one-way analysis of variance (ANOVA) tests, linear regressions, one-sided t-tests, Wilcoxon signed-rank tests, Fisher’s exact tests, and Spearman’s correlation tests in R^46^ v4.5.0. Multiple group comparisons were done for all plots in which more than two groups of data are present using ANOVA with Tukey’s HSD.

## Author Contributions

X.X. and W.R.H. conceived the study, designed the experimental and computational methods of this study, and wrote and edited the manuscript. X.X. and T.Z. collected experimental data and analyzed experimental and computational data. A.T.B. and T.F.W. implemented the computational models. J.M.C. developed code for growth rate analyses. H.F. and L.S. contributed significantly to experimental data collection. P.C.B made technical contributions and supervised part of the study. W.R.H. supervised the entire study and acquired funding.

## Acknowledgements

We specifically thank J.N.V. Martinson (Berkeley, U of MN) for the helpful feedback and for gifting us the *ΔaceA* and *ΔilvA* strains in the *Salmonella enterica* methionine-overproducer background. Gene knockout mutants in *S. enterica* 14028s were acquired from the Biodefense and Emerging Infections Research Resources Repository. We thank A. Dodge (U of MN) for training on the HPLC in L. Wackett’s group. We thank K. Yang, H. Lu, S. Klein, A. Hughes, R. Widelak, M. Soundararajan, J. Willett (U of MN), X. Wei (Oberlin College), S. Arif (UChicago), D. Singla (Vanderbilt), H. Jiao, X. Qin (XAAS), L. Lu (U of Tokyo), X. Xu, J. Jiang (Nanjing Agricultural), Q. Zhang (Xiamen Univ.), X. Liu (Nanjing Medical), P.-Y. Ho (Westlake), Y. Zhu (XJLU), X. Yi, W. Zuo (UCAS), S. Kyrazhimskiy, J. Reddan, F. She, M. Mori, T. Hwa (UCSD), J. Huisman, S. Umale, J. Gore, S. Moreno-Gámez, K. Li, L. Markey, C. Mancuso, T.D. Lieberman, Y. Yang, S. Saavedra, and R. Walton (MIT) for the helpful feedback. This work was supported by a National Institutes of Health grant (R01-GM121498) and a National Science Foundation award (No. 2425958) to W.R.H. T.Z. (2024), H.F. (2023), and L.S. (2022) were partly supported by the Undergraduate Research Opportunity Program Awards at the U of MN. X.X. was partly supported by a First-Year PhD Fellowship and an Angela Leong Fellowship through MIT.

## Competing Interest Statement

P.C.B. is a consultant to or holds equity in 10X Genomics, General Automation Lab Technologies/Isolation Bio, Next Gen Diagnostics, Cache DNA, Concerto Biosciences, Stately, Ramona Optics, Bifrost Biosystems, and Amber Bio. His laboratory has received research funding from Calico Life Sciences, Merck, and Genentech for unrelated work. All other authors disclose no competing interests.

## Data Availability

Figures were generated using Numbers v.11.2 (Mac OS) or R v4.5.0. Datasets and code are available at github.com/xion1475/antibiotic_combination_ecology.

## Notes

https://github.com/xion1475/antibiotic_combination_ecology

## Reference

1. Jackson MC, Pawar S, Woodward G. The temporal dynamics of multiple stressor effects: from individuals to ecosystems. Trends Ecol. Evol. 36, 402–410 (2021). doi: 10.1016/j.tree.2021.01.005.

2. Wu M, Xie S, Zang J, Sun Y, Xu S, Li L, Wang J. Multiple anthropogenic environmental stressors structure soil bacterial diversity and community network. Soil Biol. Biochem. 198, 109560 (2024). doi: 10.1016/j.soilbio.2024.109560.

3. Orr JA, Vinebrooke RD, Jackson MC, Kroeker KJ, Kordas RL, Mantyka-Pringle C, Van den Brink PJ, De Laender F, Stoks R, Holmstrup M, Matthaei CD, Monk WA, Penk MR, Leuzinger S, Schäfer RB, Piggott JJ. Towards a unified study of multiple stressors: divisions and common goals across research disciplines. Proc. Biol. Sci. 287(1926):20200421 (2020). doi: 10.1098/rspb.2020.0421.

4. Folt, C.L., Chen, C.Y., Moore, M.V. & Burnaford, J. Synergism and antagonism among multiple stressors. Limnol. Oceanogr. 44, 864–877 (1999). doi: 10.4319/lo.1999.44.3_part_2.0864.

5. Simmons BI, Blyth PSA, Blanchard JL, et al. Refocusing multiple stressor research around the targets and scales of ecological impacts. Nat. Ecol. Evol. 5, 1478–1489 (2021). 10.1038/s41559-021-01547-4.

6. Roemhild, R., Bollenbach, T. & Andersson, D.I. The physiology and genetics of bacterial responses to antibiotic combinations. Nat. Rev. Microbiol. 20, 478–490 (2022). 10.1038/s41579-022-00700-5.

7. Jin H, Wang L, Bernards R. Rational combinations of targeted cancer therapies: background, advances and challenges. Nat. Rev. Drug Discov. 22, 213–234 (2023). 10.1038/s41573-022-00615-z.

8. Bronstein JL, ed. Mutualism. Oxford University Press, Oxford (2015).

9. Kost C, Patil KR, Friedman J, Garcia SL, Ralser M. Metabolic exchanges are ubiquitous in natural microbial communities. Nat. Microbiol. 8(12), 2244–2252. (2023). doi: 10.1038/s41564-023-01511-x.

10. Kefford BJ, Nichols SJ, Duncan RP. The cumulative impacts of anthropogenic stressors vary markedly along environmental gradients. Glob. Chang. Biol. 29(3), 590–602 (2023). doi: 10.1111/gcb.16435.

11. Tyers M, Wright GD. Drug combinations: a strategy to extend the life of antibiotics in the 21st century. Nat. Rev. Microbiol. 17(3), 141–155 (2019). doi: 10.1038/s41579-018-0141-x.

12. Monod J. The growth of bacterial cultures. Ann. Rev. Microbiol. 3, 371–394 (1949). doi: 10.1146/annurev.mi.03.100149.002103.

13. Yeh P, Tschumi AI, Kishony R. Functional classification of drugs by properties of their pairwise interactions. Nat. Genet. 38(4), 489–494 (2006). doi: 10.1038/ng1755.

14. Brochado AR, Telzerow A, Bobonis J, Banzhaf M, Mateus A, Selkrig J, Huth E, Bassler S, Zamarreño Beas J, Zietek M, Ng N, Foerster S, Ezraty B, Py B, Barras F, Savitski MM, Bork P, Göttig S, Typas A. Species-specific activity of antibacterial drug combinations. Nature. 559(7713), 259–263 (2018). doi: 10.1038/s41586-018-0278-9.

15. Chevereau G, Bollenbach T. Systematic discovery of drug interaction mechanisms. Mol. Syst. Biol. 11(4):807 (2015). doi: 10.15252/msb.20156098.

16. Minato Y, Dawadi S, Kordus SL, Sivanandam A, Aldrich CC, Baughn AD. Mutual potentiation drives synergy between trimethoprim and sulfamethoxazole. Nat Commun. 9(1), 1003 (2018). doi: 10.1038/s41467-018-03447-x.

17. Smith, T.P., Clegg, T., Ransome, E. et al. High-throughput characterization of bacterial responses to complex mixtures of chemical pollutants. Nat. Microbiol. 9, 938–948 (2024). 10.1038/s41564-024-01626-9

18. Xiong X, Othmer HG, Harcombe WR. Emergent antibiotic persistence in a spatially structured synthetic microbial mutualism. ISME J. 18(1), wrae075 (2024). doi: 10.1093/ismejo/wrae075.

19. de Vos MGJ, Zagorski M, McNally A, et al. Interaction networks, ecological stability, and collective antibiotic tolerance in polymicrobial infections. Proc. Natl. Acad. Sci. USA114, 10666–71 (2017). 10.1073/pnas.1713372114.

20. Bottery MJ, Pitchford JW, Friman V-P. Ecology and evolution of antimicrobial resistance in bacterial communities, ISME J. 15(4) 939–948 (2021), doi:10.1038/s41396-020-00832-7.

21. Adamowicz EM, Flynn J, Hunter RC, Harcombe WR. Cross-feeding modulates antibiotic tolerance in bacterial communities. ISME J. 12(11):2723–2735 (2018). doi: 10.1038/s41396-018-0212-z.

22. Yu JSL, Correia-Melo C, Zorrilla F, Herrera-Dominguez L, Wu MY, Hartl J, Campbell K, Blasche S, Kreidl M, Egger AS, Messner CB, Demichev V, Freiwald A, Mülleder M, Howell M, Berman J, Patil KR, Alam MT, Ralser M. Microbial communities form rich extracellular metabolomes that foster metabolic interactions and promote drug tolerance. Nat. Microbiol. 7(4):542–555 (2022). doi: 10.1038/s41564-022-01072-5.

23. Klünemann, M., Andrejev, S., Blasche, S. et al. Bioaccumulation of therapeutic drugs by human gut bacteria. Nature 597, 533–538 (2021). 10.1038/s41586-021-03891-8.

24. Garcia-Santamarina S, Kuhn M, Devendran S, Maier L, Driessen M, Mateus A, Mastrorilli E, Brochado AR, Savitski MM, Patil KR, Zimmermann M, Bork P, Typas A. Emergence of community behaviors in the gut microbiota upon drug treatment. Cell. 187(22), 6346–6357.e20 (2024).

25. Crain CM, Kroeker K, Halpern BS. Interactive and cumulative effects of multiple human stressors in marine systems. Ecol. Lett. 11, 1304–1315 (2008). doi: 10.1111/j.1461-0248.2008.01253.x.

26. Côté IM, Darling ES, Brown CJ. Interactions among ecosystem stressors and their importance in conservation. Proc. R. Soc. B 283, 20152592 (2016). doi: 10.1098/rspb.2015.2592.

27. Thompson PL, MacLennan MM, Vinebrooke RD. Species interactions cause non-additive effects of multiple environmental stressors on communities. EcoSphere 9(11), e02518 (2018). doi: 10.1002/ecs2.2518.

28. Turschwell MP, Connolly SR, Schäfer RB, De Laender F, Campbell MD, Mantyka-Pringle C, Jackson MC, Kattwinkel M, Sievers M, Ashauer R, Côté IM, Connolly RM, van den Brink PJ, Brown CJ. Interactive effects of multiple stressors vary with consumer interactions, stressor dynamics and magnitude. Ecol. Lett. 25(6), 1483–1496 (2022). doi: 10.1111/ele.14013.

29. Harcombe WR. Novel cooperation experimentally evolved between species. Evolution. 64(7), 2166–72 (2010). doi: 10.1111/j.1558-5646.2010.00959.x.

30. Harcombe WR, Riehl WJ, Dukovski I, Granger BR, Betts A, Lang AH, Bonilla G, Kar A, Leiby N, Mehta P, Marx CJ, Segrè D. Metabolic resource allocation in individual microbes determines ecosystem interactions and spatial dynamics. Cell Rep. 7(4), 1104–15 (2014). doi: 10.1016/j.celrep.2014.03.070.

31. Martinson JNV, Chacón JM, Smith BA, Villarreal AR, Hunter RC, Harcombe WR. Mutualism reduces the severity of gene disruptions in predictable ways across microb ial communities. ISME J. 17(12), 2270–2278 (2023). doi: 10.1038/s41396-023-01534-6.

32. Campos AI, Zampieri M. Metabolomics-Driven Exploration of the Chemical Drug Space to Predict Combination Antimicrobial Therapies. Mol. Cell. 74(6), 1291–1303 (2019). doi: 10.1016/j.molcel.2019.04.001.

33. Xiong X, Furseth H, McCoy B, Slavic L, Zhou T, Boneza M, Yuan X, Chen S, Smanski MJ, Baughn AD, Minato Y, Harcombe WR. A duplexed CRISPRi-based approach to predict synergistic antibiotic targets in *Escherichia coli*. BioRxiv 2024.05.15.594409 (2024). doi: 10.1101/2024.05.15.594409.

34. Adamowicz EM, Harcombe WR. Weakest-Link Dynamics Predict Apparent Antibiotic Interactions in a Model Cross-Feeding Community. Antimicrob. Agents Chemother. 64(11), e00465–20 (2020). doi: 10.1128/AAC.00465-20.

35. Krueger AS, Munck C, Dantas G, Church GM, Galagan J, Lehár J, Sommer MO. Simulating Serial-Target Antibacterial Drug Synergies Using Flux Balance Analysis. PLoS One. 11(1), e0147651 (2016). doi: 10.1371/journal.pone.0147651.

36. Dukovski I, Bajić D, Chacón JM, Quintin M, Vila JCC, Sulheim S, Pacheco AR, Bernstein DB, Riehl WJ, Korolev KS, Sanchez A, Harcombe WR, Segrè D. A metabolic modeling platform for the computation of microbial ecosystems in time and space (COMETS). Nat. Protoc. 16(11), 5030–5082 (2021). doi: 10.1038/s41596-021-00593-3.

37. Douglas SM, Chubiz LM, Harcombe WR et al. Parallel mutations result in a wide range of cooperation and community consequences in a two-species bacterial consortium. PLoS One 11.e0161837 (2016). doi: 10.1371/journal.pone.0161837.

38. Bushby SRM, Hitchings GH. Trimethoprim, a sulphonamide potentiator. Br. J. Pharmac. Chemother. 33, 72–90 (1968). doi: 10.1111/j.1476-5381.1968.tb00475.x.

39. Bisesi AT, Carlson RP, Cotner L, Harcombe WR. Metabolic remodeling of microorganisms by mobile genetic elements alters mutualistic community composition. mSystems. 10(9), e0014425 (2025). doi: 10.1128/msystems.00144-25.

40. Baba T, Ara T, Hasegawa M, et al. Construction of Escherichia coli K-12 in-frame, single-gene knockout mutants: the Keio collection. Mol. Syst. Biol. 2, 0008 (2006). 10.1038/msb4100050.

41. Douglas SM, Chubiz LM, Harcombe WR, Marx CJ. Identification of the potentiating mutations and synergistic epistasis that enabled the evolution of inter-species cooperation. PLoS One, 12(5), e0174345 (2017). doi: 10.1371/journal.pone.0174345.

42. Delaney NF, Kaczmarek ME, Ward LM, et al. Development of an optimized medium, strain and high-throughput culturing methods for Methylobacterium extorquens. PLoS One 8, e62957 (2013). doi: 10.1371/journal.pone.0062957.

43. Odds FC. Synergy, antagonism, and what the chequerboard puts between them. J. Antimicrob. Chemother. 52(1),1 (2003). doi: 10.1093/jac/dkg301.

44. Ebrahim A, Lerman JA, Palsson BO, Hyduke DR. COBRApy: COnstraints-Based Reconstruction and Analysis for Python. BMC Syst. Biol. 7, 74 (2013). doi: 10.1186/1752-0509-7-74.

45. Gurobi Optimization, LLC. 2024. Gurobi Optimizer Reference Manual. gurobi.com.

46. R Core Team. R: A Language and Environment for Statistical Computing. Austria: R Foundation for Statistical Computing (2024).

