## Supplementary information for "Ecology reshapes interactions between antibiotic stressors through metabolic feedbacks"

##### **This PDF file includes:**

- Supplementary Text 1-2
- Supplementary Figures S1-S10
- Supplementary Reference

#### Supplementary Text 1: Quantitative description of the experimental microbial system

The growth physiology of *E. coli* is quite similar between the supplemented monoculture and cross-feeding conditions used throughout this paper, so comparisons between the two conditions can reveal the impact of cross-feeding. The *E. coli*-*S. enterica* community is used as a model system (Harcombe 2009; Harcombe et al., 2014) to interrogate the effect of cross-feeding on the physiology of antimicrobial responses. When grown in 200 μL drug-free Hypho minimal media in 96-well plates, *E. coli* in monoculture divides every $\sim 2.3$ hours [growth rate $\mu_{\phi}^{(Mono)} = 0.30 \text{ h}^{-1}$, $\ln(2)/0.3 = 2.31$] by plate-reader OD<sub>600</sub> (Fig. S1B-D), and the growth rate number is identical if we look at fluorescence. When 2.82 mM lactose and 80 μM methionine is provided to the monoculture *E. coli*, cells grew to a total density of $\sim 5 \times 10^8$ CFU/mL and depletes carbon first (Xiong et al., 2024).

Meanwhile, when cross-feeding with *S. enterica*, *E. coli* grows at a $\sim 77\%$ rate of its monoculture counterpart and so doubles every $\sim 3$ hours [growth rate $\mu_{\phi}^{(Co)} = 0.23 \text{ h}^{-1}$, $\ln(2)/0.23 = 3.01$]. When 2.82 mM lactose is provided to the co-culture, *E. coli* grows to $\sim 3 \times 10^8$ CFU/mL and *S. enterica* grows to $\sim 4 \times 10^7$ CFU/mL (Fig. S8H). The difference in final cell density suggests that one *S. enterica* cell can on average support the growth of 7.5 *E. coli* cells, and that the growth rates of these two coupled species are not guaranteed to be equal. Furthermore, there is excess methionine left in the media after the cross-feeding co-culture reaches the stationary phase (Fig. S1E), suggesting carbon depletion occurs first.

Growth curves shown throughout the manuscript include a long apparent lag phase ($\sim 20$ hours), which is simply a artifact of the detection limit of the plate reader. First, the detection limit of our plate reader is $\sim 220$ a.u. for CFP, $\sim 150$ a.u. for YFP, which corresponds to $\sim 0.1$ for OD<sub>600</sub>. In the current study, the starting OD<sub>600</sub> is chosen to be 0.001 per species, which is a standard small inoculum size for drug susceptibility tests to get rid of spontaneous mutants. With a detection limit of OD<sub>600</sub> = 0.1, cells have to divide $\sim 6.6$ times before reaching the detection limit, and this gives an artificial lag time of 15–19 hours (Fig. S1D).

#### Supplementary Text 2: A genome-scale metabolic model predict that the change in antibiotic interaction is driven by metabolism

We modeled the impact of perturbing 40 gene targets in monoculture and co-culture (Materials & Methods). The simulated drug targets were not chosen to match our experimental conditions but rather to provide an unbiased distribution of effects. For each targeted reaction, we reduced the stoichiometric coefficients of all metabolic products to a fraction ($1/\alpha$) of its unperturbed counterpart and diverted the remaining metabolic substrates to unreactive waste (Fig. S6A and Materials & Methods; Krueger et al., 2016). For 40 metabolic targets spanning 24 metabolic pathways (Fig. S6B), we identified values of $\alpha$ that reduces growth of *E. coli* and *S. enterica* in monoculture by 50% in the flux balance analysis (FBA). Using dynamic FBA (dFBA) in COMETS, we incorporated these unique $\alpha$ values to simulate growth in monoculture and co-culture when each of the 40 reactions was targeted singly and in all 780 pairwise combinations.

The simulations supported that drug combinations will often have different impacts in monoculture than in cross-feeding co-culture. In the majority of pairs (640/780, 83%) we observed that $X$ scores differed for *E. coli* by more than 10% between cross-feeding co-culture and monoculture. We observed a change in the class of drug interaction 52% of the time. In contrast to our experimental results, we observed that $X$ scores increase roughly as often as they decrease (Fig. S6C). Consistent with previous research (Bronchado et al., 2018), stressors targeting the same pathway tend to produce synergistic interactions, while stressors targeting multiple pathways tend to produce antagonistic ones (Fig. S6D).

We found that growth rate alone are rarely the primary mechanism through which cross-feeding alters the effect of stressor combinations (Fig. S7A). For each gene perturbation pair, we calculated the average difference in single-drug growth rates between monoculture and cross-feeding co-culture. Difference in raw rates for *E. coli* and *S. enterica* are $\Delta\mu_E^{single}$ and $\Delta\mu_S^{single}$, respectively. Difference in relative rates for *E. coli* and *S. enterica* are $\Delta g_E^{single}$ and $\Delta g_S^{single}$, respectively. We observed that at a given growth rate difference, there are almost always a diverse distribution of differences in the $X$ scores.

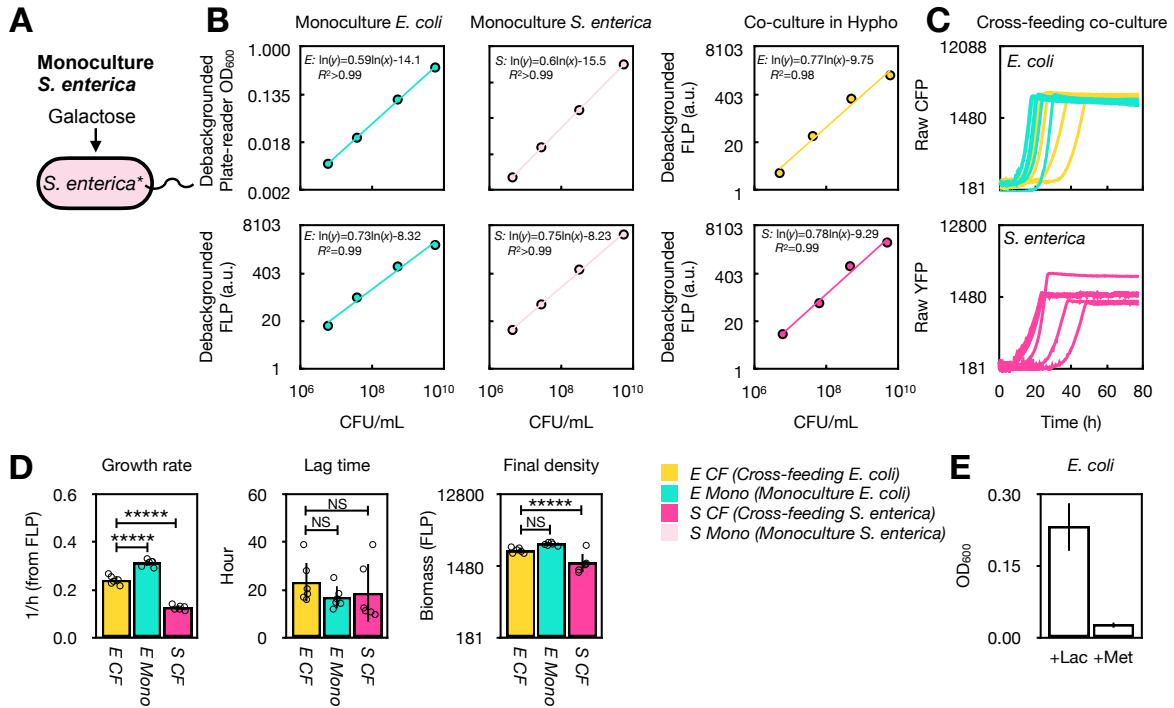

**Figure S1. Genome-integrated fluorescent proteins enable species-specific measurement of growth rates and abundance in the cross-feeding co-culture.** **A** *S. enterica* monoculture is supplemented with galactose in Hypho minimal media (Materials & Methods). **B** CFP (cyan fluorescent protein) and YFP (yellow fluorescent protein) signals correlate with species-specific CFU/mL counts ($R^2 > 0.93$, consistent with Adamowicz et al., 2018). *E. coli* and *S. enterica* monocultures are grown to the stationary phase in test tubes with 200-rpm shaking at 37°C, and serially diluted in 96-well plates containing species-specific Hypho media. CFU and fluorescence are measured for each dilution. A mixed (artificial) co-culture is created by combining concentrated monocultures at a 1:1 volume ratio and serially diluting in Hypho lactose media. Slopes of linear regression are equal across species with no cross-fluorescence, allowing for direct comparisons of growth rates across species. **C** Cross-feeding *E. coli* exhibits similar growth dynamics with the monoculture counterpart. Three replicates have been shown in Fig. S1C in a previous manuscript (Xiong et al., 2024); here we included more independent replicates. **D** Quantification of growth parameters from left to right: growth rates, apparent lag times, and final yields. They are estimated by fitting log-linear models to species-specific fluorescent growth curves. In co-culture, *E. coli* grows at ~77% of its monoculture growth rate (one-way ANOVA with Tukey's HSD, $P < 1 \times 10^{-6}$), with no significant differences in apparent lag time or yield. Consistent with Harcombe et al. (2014), Adamowicz et al. (2018), and Xiong et al. (2024), *E. coli* outgrows *S. enterica* when cross-feeding in growth rate and final density (both $P < 1 \times 10^{-6}$, one-way ANOVA with Tukey's HSD). Bars show means ± 1 SD away from mean. Open circles represent single data points. When more than two groups of data are shown in the same plot, multiple-test comparison corrections were done and $P$-values were adjusted. Statistical significance annotations: NS: $P \geq 0.1$; \$: $0.05 \leq P < 0.1$; \*: $10^{-2} \leq P < 0.05$; \*\*: $10^{-3} \leq P < 10^{-2}$; \*\*\*: $10^{-4} \leq P < 10^{-3}$; \*\*\*\*: $10^{-5} \leq P < 10^{-4}$; \*\*\*\*\*: $P < 10^{-6}$. These conventions are used throughout the manuscript. **E** The two-species community enters the stationary phase because lactose runs out. In a two-day stationary phase culture, spent media were collected by filtering out cells and supplemented with 2.82 mM lactose or 80 μM methionine, respectively. Log-phase *E. coli* was inoculated into the supplemented media. Only the spent media with lactose supplementation shows robust growth.

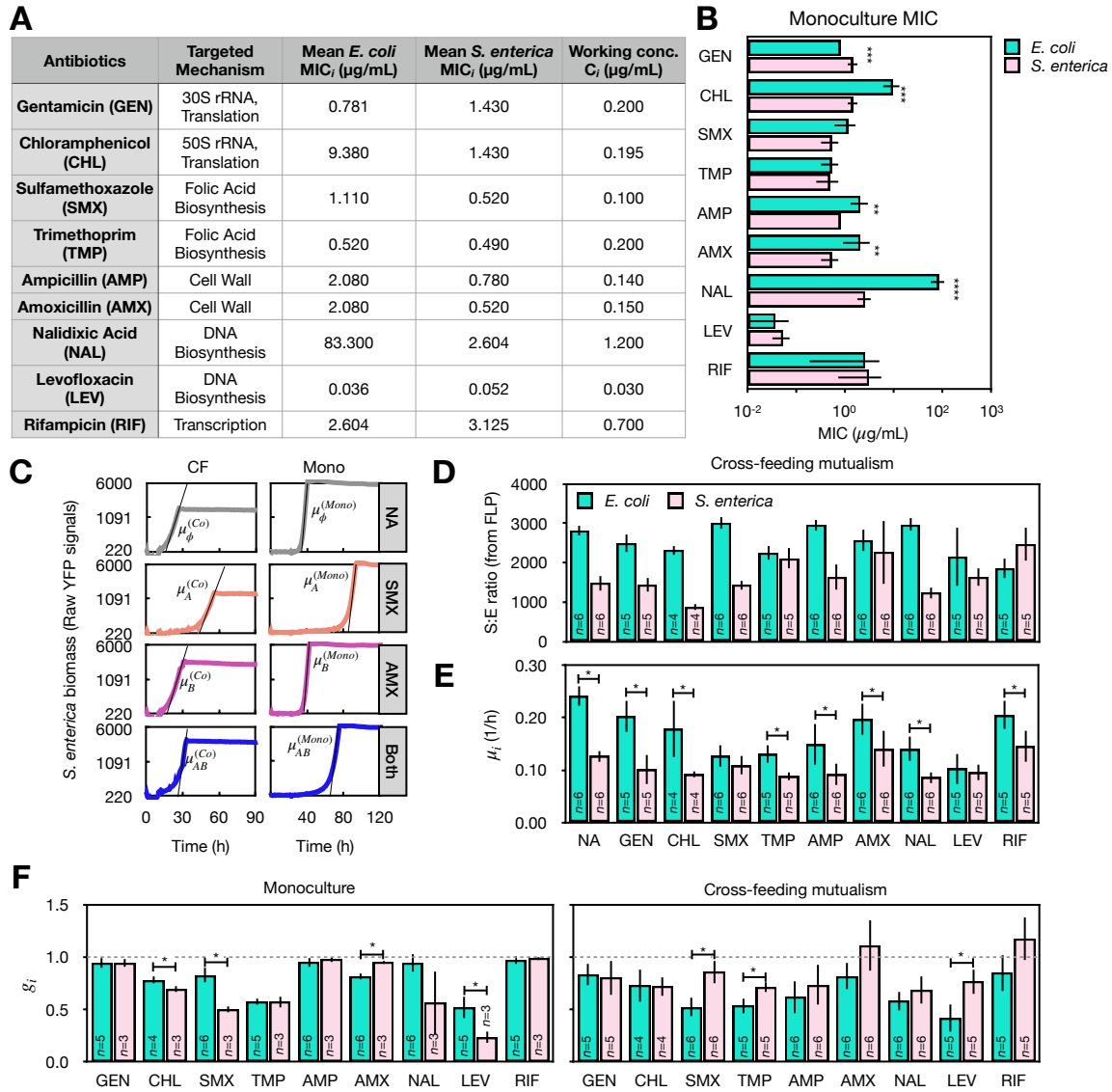

**Figure S2. Relative growth rates ($g_i$) in *E. coli* range between 60% and 95% of their no-drug counterpart in monoculture and cross-feeding co-culture in 9 diverse antibiotics at their sub-MIC working concentrations ($C_i$). A-B** Minimum inhibitory concentrations (MICs) of *E. coli* and *S. enterica* were measured in their respective monocultures for each of the 9 drugs as the lowest concentration necessary to fully inhibit growth (OD<sub>600</sub> is <10% of the no-drug counterpart, i.e. MIC<sub>90</sub>) after 48 hours. Average MIC<sub>i</sub> values for each drug $i$ are determined from six biological replicates ($n = 6$) across two independent experiments. The working concentration ($C_i$) for each drug $i$ is selected as the lowest sub-MIC concentration that reduces growth yields by at least 10% after 24 hours compared to the no-drug control in a previous experiment. **C** Representative growth curves and growth rate ($\mu_i$) estimations of *S. enterica* are shown as in Fig. 1B. **D** Final densities of *E. coli* and *S. enterica*, measured by FLP signals at the end of incubation under single-antibiotic or no-antibiotic (NA) conditions in cross-feeding co-cultures. **E** Raw growth rates ($\mu_i$) of *E. coli* and *S. enterica* in cross-feeding co-cultures under single-antibiotic or no-antibiotic conditions. **F** Relative growth rates ($g_i$) of *E. coli* and *S. enterica* under each single antibiotic at their respective working concentrations (Fig. S2A) in monocultures and cross-feeding mutualism. To ensure that every pairwise antibiotic combination is assayed in three independent experiments, each single drug is concurrently tested in each experiment.

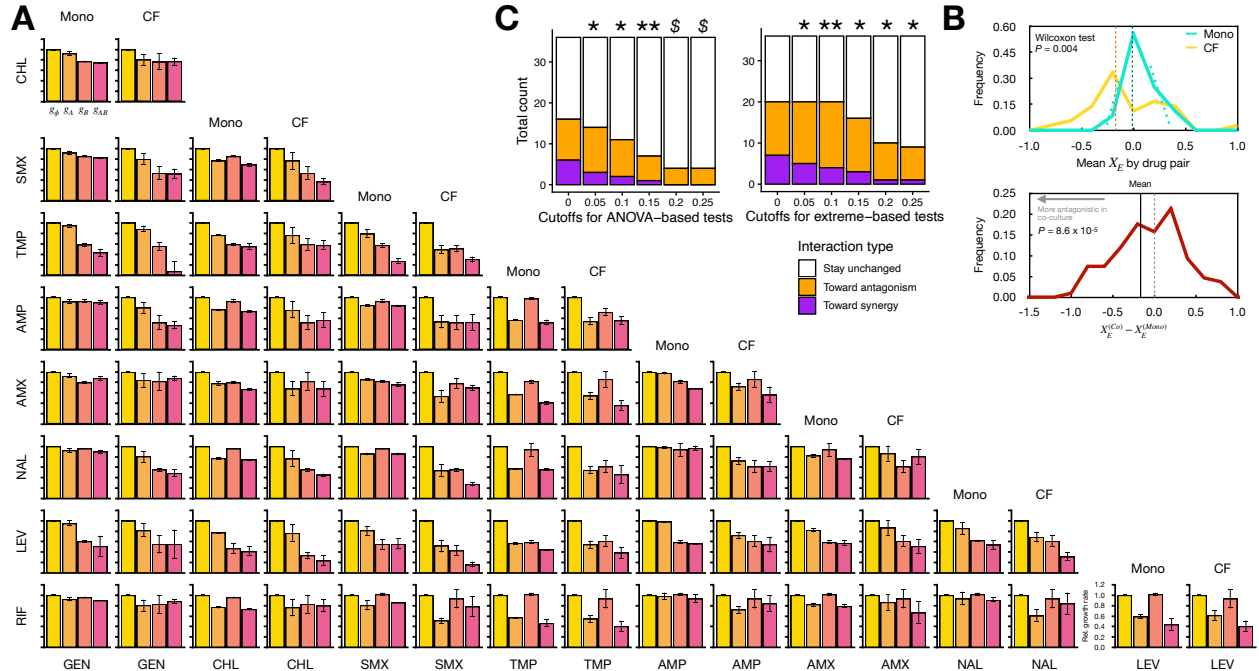

**Figure S3. Interaction indices ($X$) of antibiotic pairs generally become smaller for cross-feeding *E. coli*. A**

Relative growth rates for monoculture (Mono) and cross-feeding (Mutual) *E. coli* without drugs ($g_\phi$), under antibiotics individually ($g_A, g_B$) and in pairs ($g_{AB}$). All data shown are from three independent experiments. Note that although *E. coli* grown in single antibiotics have been assayed up to six independent times (Fig. S2D-F), only three of the values that were collected in the same experiments with the antibiotic combinations appear here for rigorous comparisons. **B** (Top) Mean antibiotic interaction indices ($X$) for monoculture and cross-feeding *E. coli* plotted by individual drug pairs. Monoculture $X$ scores have a mean of 0, with its peak centered at 0 and dispersed across an SD = 0.10 (dotted cyan peaks), but the co-culture $X$ is on average smaller. Vertical dotted lines denote average $X$ scores in either ecological conditions. (Bottom) Differences in $X$ scores between co-culture and monoculture *E. coli* is on average negative across all pairwise comparisons ($n = 108$). $P$-value is calculated by comparing the mean of the population against 0 with a one-way ANOVA test. **C** More drug pairs shift toward antagonism for *E. coli* from monoculture to cross-feeding co-culture than toward synergy. This result is robust regardless of cutoffs of $X$ for classifying an interaction as synergy or antagonism, and regardless of whether the interaction is classified using an ANOVA test against the cutoff $X$ (left) or using extreme values (right) of $X$ against the cutoff as we chose for Fig. 1D. Most drug interaction types remain unchanged. Binomial test is performed for each classification method and at each classification cutoff $X$ between numbers of drug pairs shifting toward antagonism and toward synergy; \* denotes $P < 0.05$ and \$ denotes $P < 0.1$.

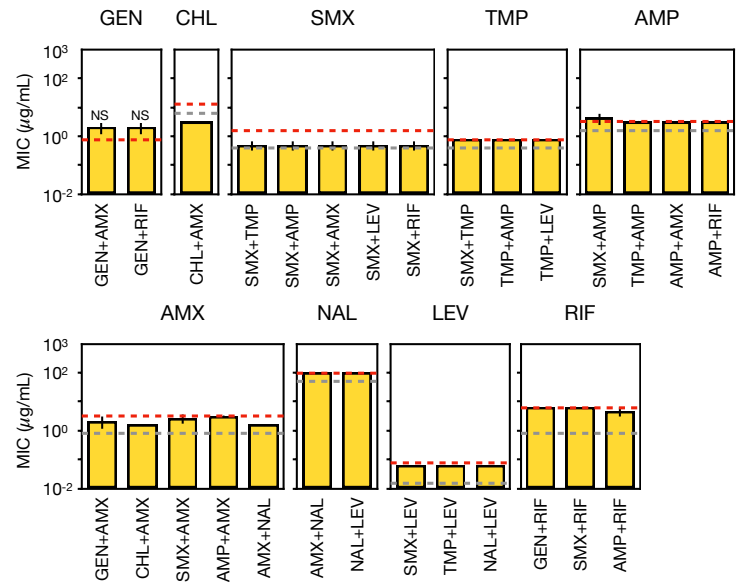

**Figure S4. Cross-feeding *E. coli* grown with antibiotic combinations does not evolve higher MICs.** We ruled out the hypothesis that higher antibiotic resistance evolved during growth and underlies the shift towards antibiotic antagonism/away from synergy for cross-feeding *E. coli*. In the eleven relevant co-cultures, resistance to each single drug was assayed at growth saturation (Fig. 1D) by measuring MICs according to our previously developed protocol (Xiong et al., 2024). Briefly, the stationary co-cultures from the screen experiment (Fig. 1C and S3) were diluted to $OD_{600} = 0.002$ into fresh Hypho media with lactose and methionine. In this environment, *E. coli* can independently grow without relying on *S. enterica*, while *S. enterica* still relies on *E. coli*. Thus, given the weakest link hypothesis (Adamowicz et al., 2018), even if *S. enterica* has higher MIC than *E. coli* for some antibiotics, there will not be no community growth and the MIC measurement from these cultures should still reflect *E. coli*'s MICs (Xiong et al., 2024). We found that the antibiotic resistance levels in cross-feeding *E. coli* are within the range from ancestral colonies (Fig. S2A-B). Red and grey dashed lines denote highest and lowest MIC levels measured from freshly streaked *E. coli* colonies, respectively.

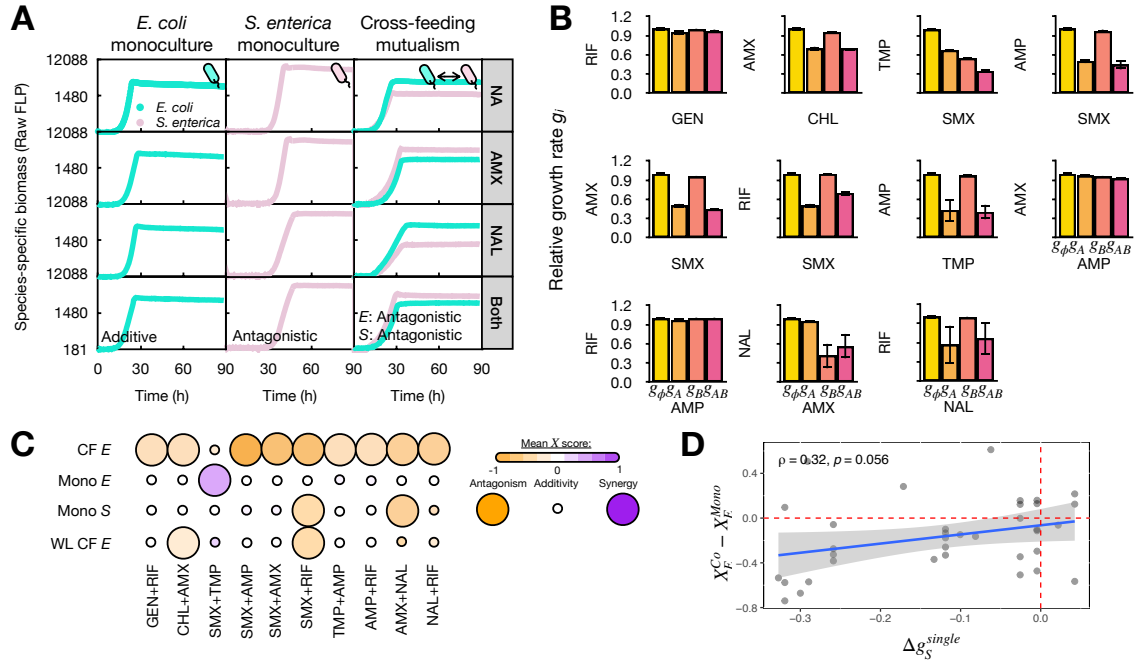

**Figure S5. The weakest-link hypothesis can only explain a fraction of cases where antibiotic interactions are classified differently between monoculture and cross-feeding *E. coli*.** **A** One representative set of growth curves of *E. coli* and *S. enterica* from the screen experiment in the antibiotic combination of AMX and NAL. **B** The growth rates of monoculture *S. enterica* were separately measured in the same format as in Fig. 1B and S2C in all eleven drug pairs shifting toward antagonism in co-culture *E. coli*. **C** Antibiotic interaction types for monoculture *S. enterica* (Mono *S*) and predicted drug interactions for cross-feeding *E. coli* (WL CF *E*) following the weakest link (WL) hypothesis. Corresponding data for *E. coli* (monoculture: Mono *E*; cross-feeding: CF *E*) were reproduced from Fig. 1D for antibiotic pairs that shift toward antagonism and away from synergy in cross-feeding *E. coli*. The weakest link (WL) classification was predicted from both monoculture growth rates. We defined the “weakest-link” effect here as the notion that when drugs are present in the cross-feeding co-culture, each species will adopt the slower relative growth rate ($g_i$) of the monocultures. For these relevant drug pairs, the “weakest-link” hypothesis only predicts two drug interactions correctly for cross-feeding *E. coli*. The drug interaction profiles in monoculture *S. enterica* predict correctly in two drug interactions. Legend is identical to that in Fig. 1D. **D** Correlation between mean relative growth rate differences in single-drug conditions and mean differences in $X$ between monoculture and co-culture in the experiment in Fig. S3A.

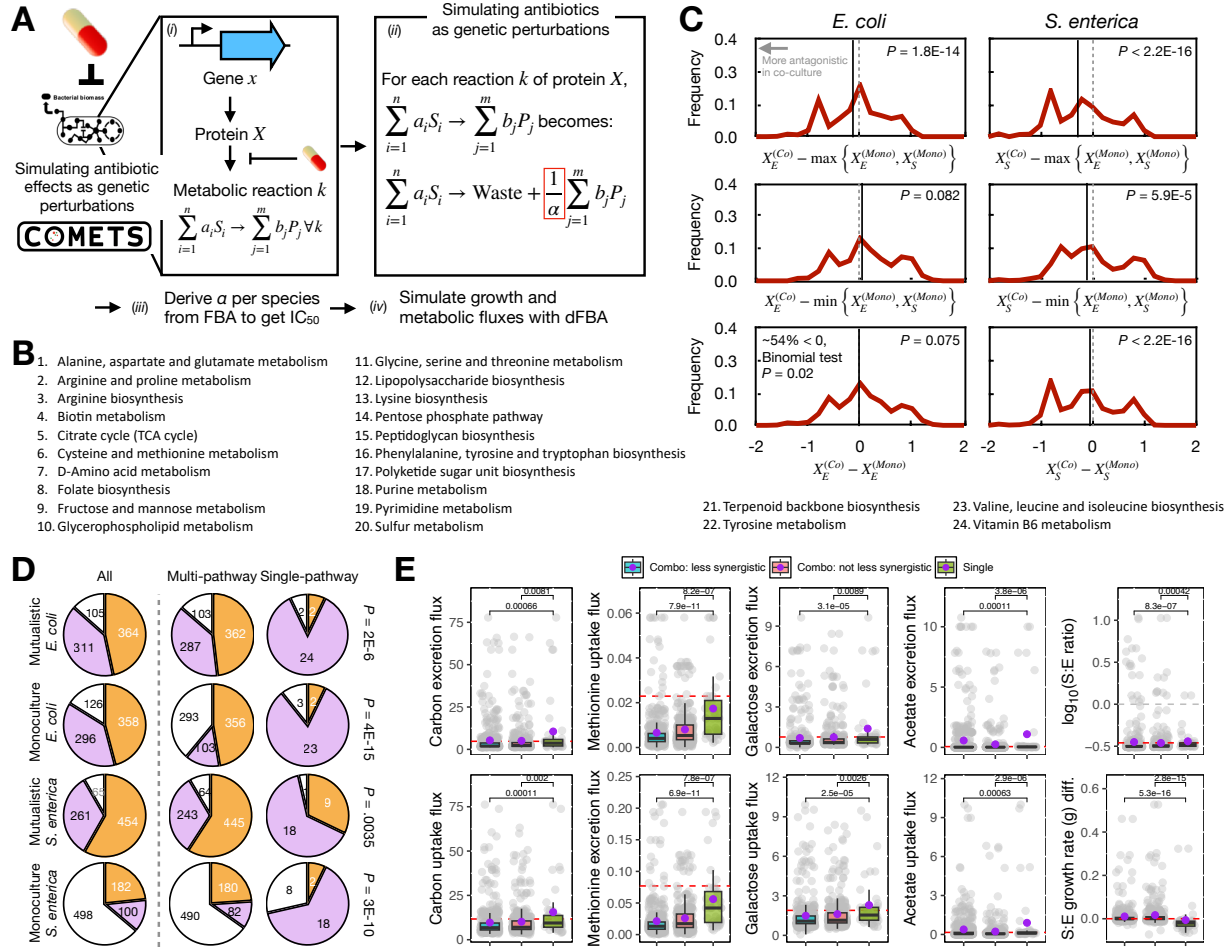

**Figure S6. A genome-scale metabolic model also predicts shifts toward antagonistic stressor interactions in cross-feeding *E. coli*.** **A** Schematic of our simulation procedure. Step (i): We simulated the physiological effect of pairwise genetic perturbations on all metabolic fluxes that a target gene controls, following an established flux diversion method in Kreuger et al. (2016). Step (ii): For an antibiotic targeting gene $x$, all metabolic reactions ($k$) for which protein $X$ is responsible experience a smaller flux where the chemical products are reduce to a fraction ($1/\alpha$) of the non-perturbed state such that the growth rate measured in flux balance analysis (FBA) is reduced by half, and the remaining products are considered non-reactive waste. Step (iii): We measured $\alpha$ for each species separately as the value that would reduce growth by exact half ($IC_{50}$) in FBA. If the gene is not essential (i.e. no growth rate reduction in FBA), then $\alpha$ was set to 100,000. Step (iv): We ran dynamic FBA (dFBA) in the COMETS platform (Dukovski et al., 2021) to simulate complete growth curves for *E. coli* in both cross-feeding and monoculture contexts, under conditions with no gene perturbations, single-gene perturbations, and gene-pair perturbations. We used the same metric, $X$, to measure the interaction from perturbing gene pairs. **B** All 24 metabolic pathways that the 40 perturbed genes are involved with. The KEGG pathways are identified by COMETS. **C** A substantial fraction of pairwise perturbations results in reduced synergy and cannot be explained by either monoculture effects. Distribution plots for differences in the antibiotic interaction indices ($X$) between co-culture and monocultures are shown as in Fig. S3B. **D** Distribution of antibiotic interaction types for *E. coli* and *S. enterica* in monoculture and the cross-feeding co-cultures (All, left). Perturbation pairs involved in the single (identical) pathways are significantly more enriched in synergistic interactions (Fisher's exact tests, $P < 6 \times 10^{-11}$). $X$ scores for cross-feeding

*E. coli* is more often smaller than the monoculture counterpart (Binomial test, $P = 0.02$). **E** Pairwise genetic perturbations result in different metabolic exchange and growth rates in both species than single-gene perturbations do. Metabolic exchange in the cross-feeding co-culture was measured as the maximum excretion or uptake flux (mmol/gDW/hour) within 20 hours ($\pm 10$) around the mid-log point of the growth curve in each species, respectively. Each dot represents one perturbation condition (i.e. one gene-pair perturbation, or one single-gene perturbation). The S:E ratio refers to the species ratio between *S. enterica* and *E. coli* yields at the end of growth for all co-cultures. Grey dashed line denotes a 1:1 ratio. The S:E growth rate (*g*) difference (diff.) indicates the relative *E. coli* growth rates subtracted from the *S. enterica* counterpart. Higher difference indicates more growth rate advantages for *S. enterica*, where *E. coli* usually grows faster in the no-drug mutualism. Statistical comparisons were performed with pairwise Wilcoxon tests and adjusted *P*-values were shown. Red dashed lines denote levels in each plot for the unperturbed control. Each purple dot denotes the average value of each group.

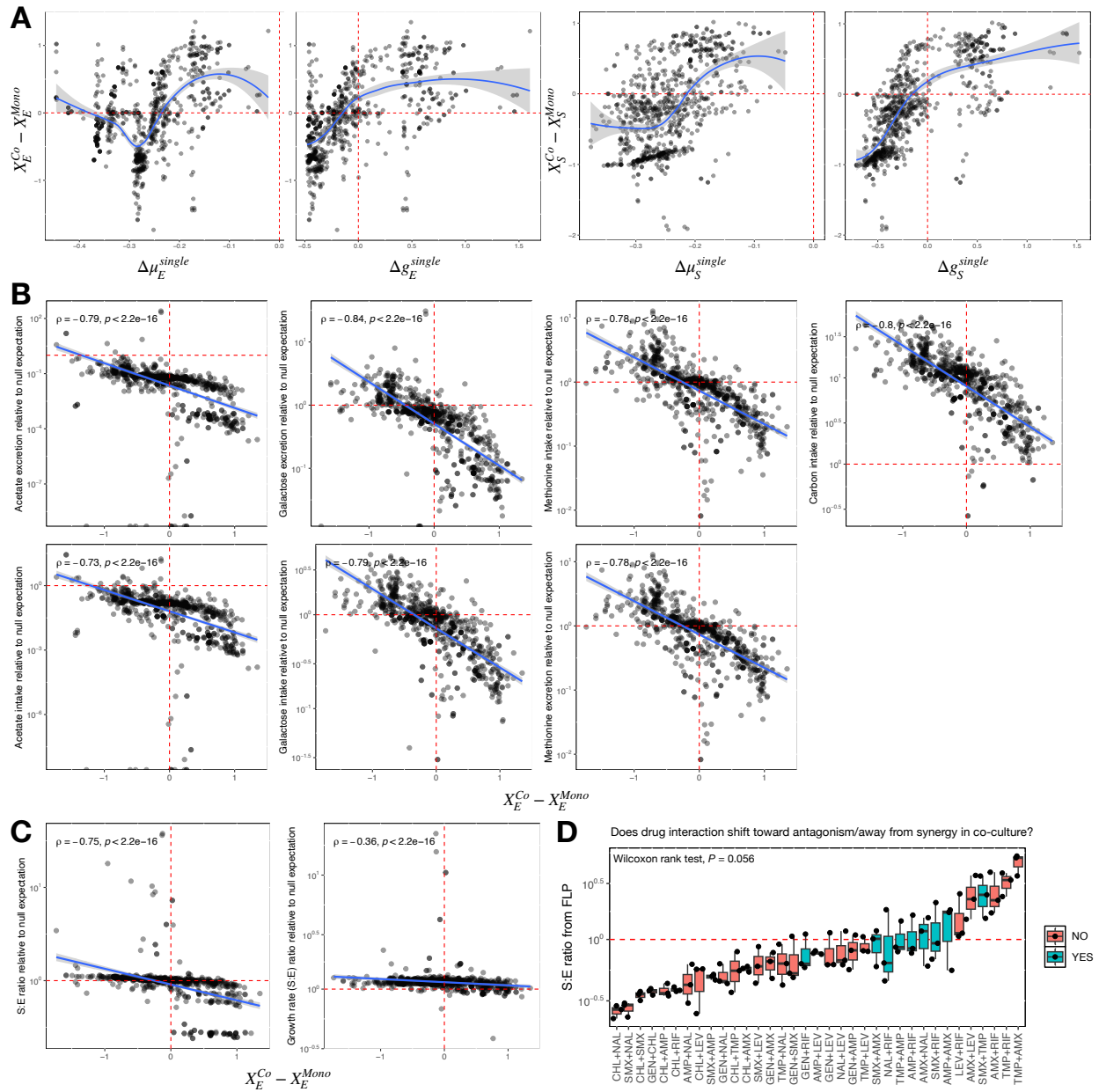

**Figure S7. Non-additive effects of pairwise gene perturbation on metabolic excretion, not growth rate changes alone, affect shifts in stressor interactions for cross-feeding *E. coli*.** **A** Correlation between mean growth rate differences in single-gene conditions and differences in stressor interaction indices ($X$) between monoculture and co-culture in simulations. Blue lines are the Loess lines of fit. **B-C** Changes in stressor interaction indices ($X$) for *E. coli* are correlated with metabolic fluxes in dual perturbation conditions relative to the linear (null) expectation from both involved single perturbations. Fluxes measured from Fig. S6E for *E. coli* (top) and *S. enterica* (bottom), respectively. Vertical lines denote no change in stressor interaction; horizontal lines denote matched linear expectations. **C** Species ratios and relative growth rate ratios are also correlated with changes in $X$. **D** Experimental S:E ratios from YFP:CFP signals across all stationary antibiotic pair conditions in the cross-feeding co-culture. For each drug pair, distribution of biological replicates is shown with the middle line denoting median. Red dashed line indicates equal species frequency.

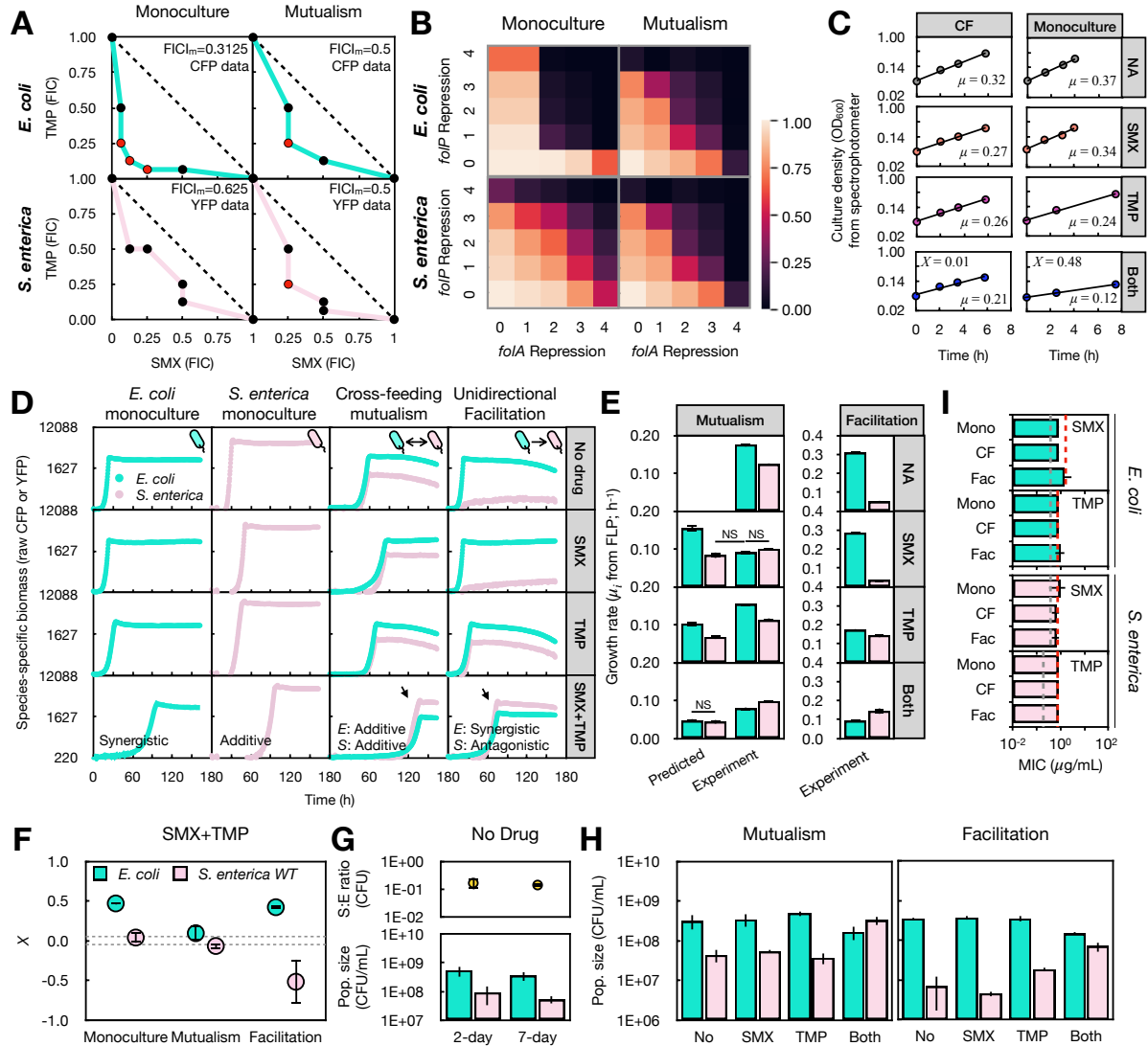

**Figure S8. Checkerboard assays, the genome-scale metabolic model, and drug exposure experiments in log-phase test tube cultures confirm the weakened SMX-TMP synergy for cross-feeding *E. coli*.** A Checkerboard assays (see Methods) recapitulate the reduced SMX-TMP synergy in cross-feeding (Mutualism, borderline synergy: FIC<sub>m</sub> = 0.5) *E. coli* compared to its monoculture counterpart (Monoculture, synergy: FIC<sub>m</sub> = 0.3125). Data from one representative experiment is shown from two independent ones. Red dots denote antibiotic concentration pairs with FICI ≤ 0.5. B COMETS similarly predicts the reduced synergy in cross-feeding *E. coli*. Checkerboard assays are simulated at four different levels (1~4) of perturbation in the metabolic fluxes that each target gene is involved in. SMX targets *folP*, and TMP targets *folA*. Colors in each box denote simulated growth rates with respect to the no-perturbation control (bottom left corner boxes). *E. coli* has faster relative growth in certain perturbation combinations when cross-feeding than in monoculture. C Reduced SMX-TMP synergy in cross-feeding co-cultures does not require lag-phase biology. OD<sub>600</sub> was measured with the spectrophotometer module on a SpectraMax M Series Multi-Mode Microplate Reader (Molecular Devices, CA). Monocultures of *E. coli* and cross-feeding co-cultures of *E. coli* and *S. enterica* (CF) were pre-grown to mid-log phase (OD<sub>600</sub> = 0.2) separately, and 5 mL culture was added into 30-mL test tubes at starting OD<sub>600</sub>=0.05. Drugs were then added at working concentrations (Fig. S2A), and tubes were incubated at 37°C with shaking. An amount of 750 μL culture was sacrificed in each

measurement. Black lines are lines of best fit to the growth curve data (shown in colored bubbles), and growth rates ($\mu_i$) are slopes of these fit lines on a natural log-scaled OD<sub>600</sub>. Consistent with in 96-well plates, the cross-feeding co-culture grows more slowly than the *E. coli* monoculture without drugs. Drugs at our concentrations only reduce growth rates but do not increase lag time. The SMX-TMP synergy also disappears in the cross-feeding co-culture (see *X* scores). **D** Representative CFP (cyan, *E. coli*) and YFP (pink, *S. enterica*) growth curves in the SMX-TMP combination among three more repetitions (one repetition is shown). In the unidirectional facilitative co-cultures and *E. coli* monocultures, lactose and methionine are supplemented so that *S. enterica* still depends on *E. coli* but not *vice versa*. Arrows indicate species dominance switches. **E** *S. enterica* in the cross-feeding co-culture does not grow faster ($\mu_i$) than *E. coli* under no antibiotics, SMX alone, and TMP alone. “Predicted” values are estimated by multiplying the relative growth rates (*g*) of each species in their monocultures with SMX alone and TMP alone, respectively, with the raw growth rates of each species in the respective drug-free co-culture. *E. coli* is predicted to grow faster in both single-antibiotic conditions. For clear view of data, pairs of growth rates **not** significantly different are labeled (NS) and shown, all other pairwise comparisons in each plot yield significant differences. **F** Antibiotic interaction indices (*X*) measured from each ecological condition with the new biological triplicates. Dashed lines are as shown in Fig. 1C. **G** The S:E ratio from CFU/ml counts are stable in Hypho media for a week. All CFU counts are measured on day 7 in this study to allow sufficient time for the antibiotic-treated communities to reach the stationary phase, while drug-free communities usually do so by day 2 or 3. Both species die a little from day 2 to day 7, but at a similar rate. **H** Raw CFU/mL counts from selective plating of *E. coli* and *S. enterica* in the cross-feeding (Mutualism) and facilitative (Facilitation) co-culture. **I** *E. coli* and *S. enterica* have not evolved higher resistance from the end of growth in both SMX and TMP. MIC against either drug is measured as in Fig. S4. Mono: monoculture; CF: cross-feeding mutualism; Fac: facilitative co-culture.

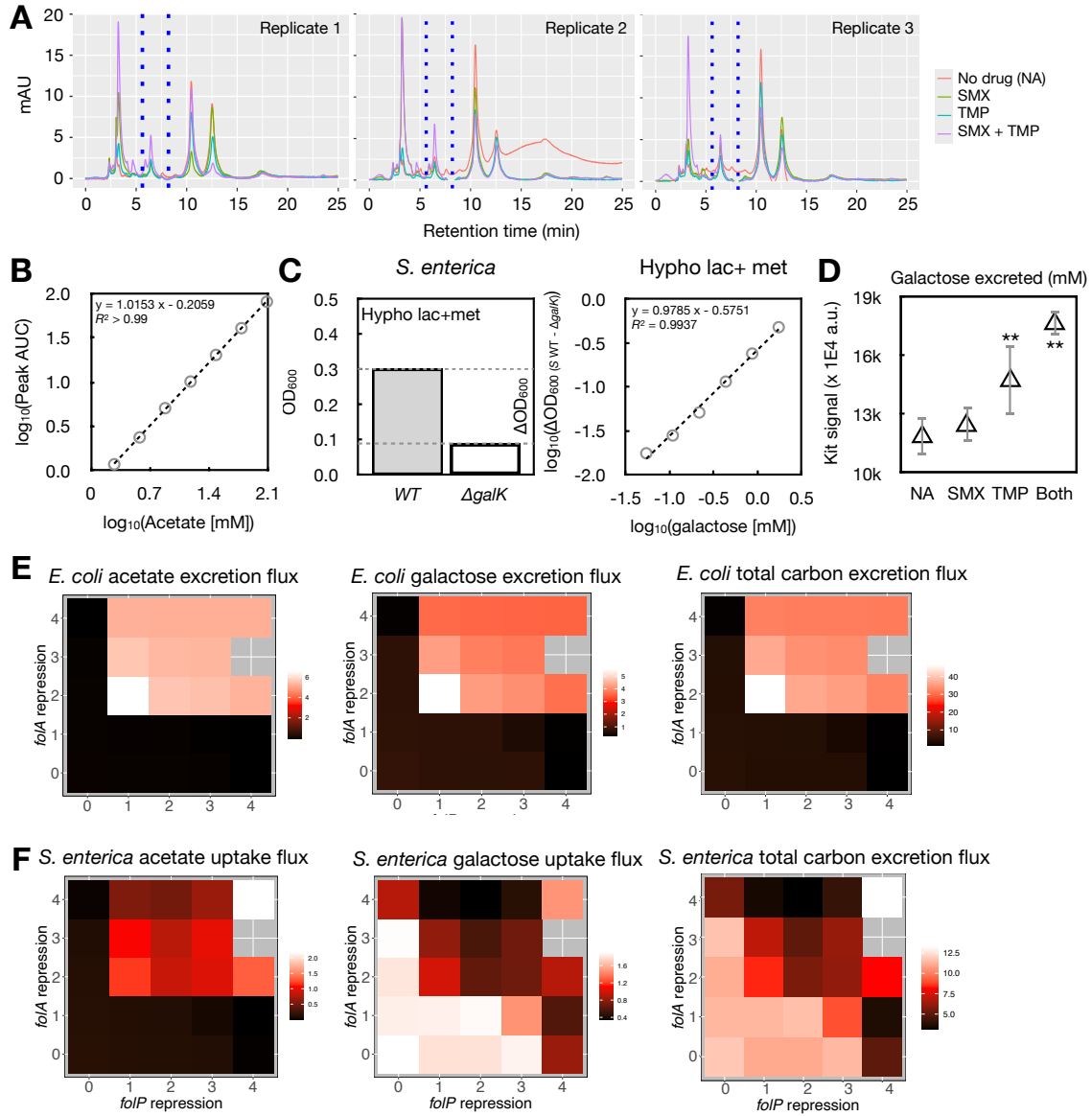

**Figure S9. Spent media analysis and the genome-scale metabolic model agree on the *E. coli* carbon over-excretion when perturbed by both SMX and TMP.** **A** Raw peak plots in HPLC experiments across triplicates of *E. coli* monocultures. Acetate peak: 7~9 min (blue dashed lines). **B** The standard curve between acetate concentration and the area under the curve (AUC) of the chemical peak. The peak before 5 min is caused by metals in the Hypho media. **C** Method for measuring galactose excretion in *E. coli*. (Left) The difference in growth yields ($\Delta OD_{600}$) between *S. enterica* WT and $\Delta galK$ strains grown in Hypho media with lactose (lac), methionine (met), and galactose allows measurement of galactose concentration. (Right) A standard curve shows that $\Delta OD_{600}$ between the two strains is highly correlated with galactose supplementation. **D** Measurement of galactose signal strengths in the spent media collected strictly at $OD_{600} = 0.21$ using the Amplex Red Galactose/Galactose Oxidase Assay Kit (Thermo Fisher, MA). **E-F** COMETS simulations agree with experiments on carbon excretion in *E. coli*, which is sum of all carbon in galactose and acetate. Carbon excretion (*E. coli*) and uptake (*S. enterica*) fluxes (unit: mmol/gDW/h) are derived from the growth simulations in Fig. S6.

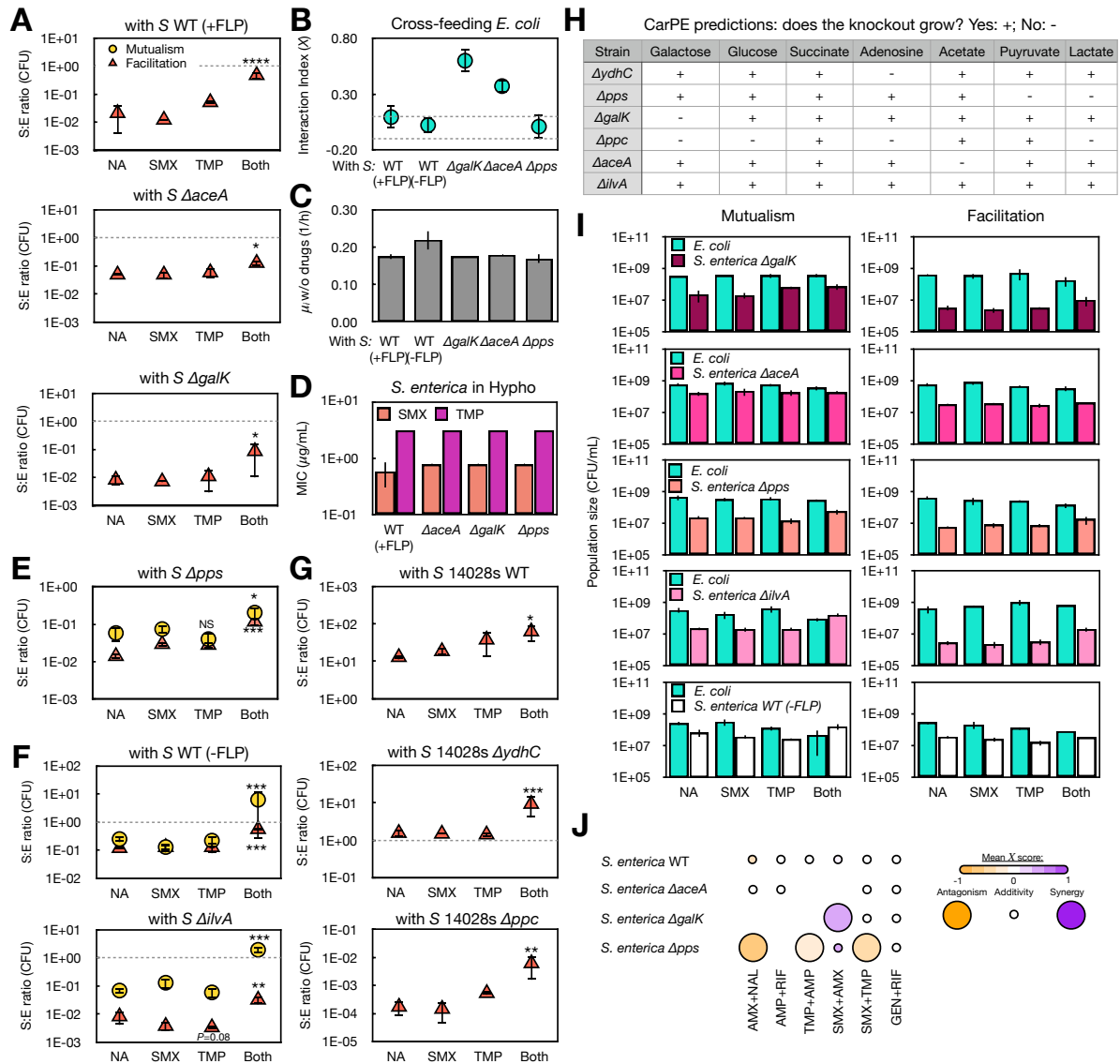

**Figure S10. *E. coli* experiences more synergistic antibiotic interactions (higher $X$) when cross-feeding with *S. enterica* $\Delta aceA$ or $\Delta galK$ mutants against SMX-TMP as well as other drug pairs. **A** S:E ratios from CFU/mL data for unidirectionally facilitative co-cultures when *E. coli* grows with *S. enterica* WT, $\Delta aceA$, or $\Delta galK$ strains under different drug conditions. **B** Interaction indices ($X$) for *E. coli* that cross-feeds with different *S. enterica* strains. Cross-feeding with *S. enterica* WT strains with and without the fluorescent gene has minimal effects on the SMX-TMP interaction for *E. coli*. Dashed lines are as shown in Fig. 1C. **C** *E. coli* does not experience a growth defect when cross-feeding with different *S. enterica* strains ($n=3$). **D** Deletion of $\Delta galK$, $\Delta aceA$, or $\Delta pps$ does not change resistance to either SMX or TMP in *S. enterica*. MIC for *S. enterica* $\Delta galK$ strain is measured in monoculture Hypo glucose media, and the rest in Hypo galactose media. Mutants are constructed using the non-fluorescent *S. enterica* background. **E** S:E ratios from CFU/mL data for mutualistic and facilitative co-cultures between *E. coli* and various *S. enterica* strain under different drug conditions. Dashed lines indicate equal *E. coli* and *S. enterica* CFU/mL counts (same below). **F-G** No evidence that the SMX-TMP pair induces *E. coli* to excrete lactate, pyruvate, adenosine, or glucose. **(F)** Under both SMX and TMP, *S. enterica* (*S*) $\Delta ilvA$ and the non-fluorescent WT strains both grow to higher final fractions when cross-feeding with *E. coli*. **(G)** When grown in a facilitative co-culture with *E.***

*coli*, the non-methionine-producing *S. enterica* 14028S $\Delta ydhC$ and $\Delta ppc$ mutants also grow to higher fractions under both SMX and TMP, suggesting that *E. coli* likely does not excrete more adenosine (*ydhC*) or other carbon sources (*ppc*) under the drug combination. **H** Predicted presence (Yes: +) and absence of growth (No: -) in *S. enterica* mutants based on the *E. coli* counterparts grown in single carbon sources in M9 media (CarPE, Tong et al., 2020). **I** Raw CFU counts for the cross-feeding (Mutualism) and facilitative (Facilitation) co-cultures between *E. coli* and various non-14028S *S. enterica* strains. **J** Antibiotic interaction types for monoculture *S. enterica* $\Delta aceA$, $\Delta galK$, and $\Delta pps$ strains under six important drug pairs. Measurement approaches are identical as in Fig. 1B except that the growth is measured by tracking OD<sub>600</sub> over time for monoculture *S. enterica*.

### Supplementary Reference

- Adamowicz EM, et al. 2020. Cross-feeding modulates the rate and mechanism of antibiotic resistance evolution in a model microbial community of *Escherichia coli* and *Salmonella enterica*. *PLoS Pathog.* 16(7), e1008700.
- Brochado AR, et al. 2018. Species-specific activity of antibacterial drug combinations. *Nature* 559, 259-263.
- Campos AI, Zampieri M. 2019. Metabolomics-driven exploration of the chemical drug space to predict combination antimicrobial therapies. *Mol Cell.* 74(6), 1291-1303.
- Demidenko E, Miller TW. 2019. Statistical determination of synergy based on Bliss definition of drugs independence. *PLoS One.* 14(11), e0224137.
- Krueger AS, et al. 2016. Simulating serial-target antibacterial drug synergies using flux balance analysis. *PLoS One.* 11(1), e0147651.
- Minato Y, et al. 2018. Mutual potentiation drives synergy between trimethoprim and sulfamethoxazole. *Nat Commun.* 9(1), 1003.
- Tong M, et al. 2020. Gene Dispensability in *Escherichia coli* Grown in Thirty Different Carbon Environments. *mBio.* 11(5), e02259-20.
- Xiong X, et al. 2024. Emergent antibiotic persistence in a spatially structured synthetic microbial mutualism. *ISME J.* 18(1), wrae075.
